# Introgressive and horizontal acquisition of *Wolbachia* by *Drosophila yakuba*-clade hosts and horizontal transfer of incompatibility loci between distantly related *Wolbachia*

**DOI:** 10.1101/551036

**Authors:** Brandon S. Cooper, Dan Vanderpool, William R. Conner, Daniel R. Matute, Michael Turelli

## Abstract

Maternally transmitted *Wolbachia* infect about half of insect species, yet the predominant mode(s) of *Wolbachia* acquisition remains uncertain. Species-specific associations could be old, with *Wolbachia* and hosts co-diversifying (i.e., cladogenic acquisition), or relatively young and acquired by horizontal transfer or introgression. The three *Drosophila yakuba*-clade hosts ((*D. santomea, D. yakuba*), *D. teissieri*) diverged about three million years ago and currently hybridize on Bioko and São Tomé, west African islands. Each species is polymorphic for nearly identical *Wolbachia* that cause weak cytoplasmic incompatibility (CI)–reduced egg hatch when uninfected females mate with infected males. *D. yakuba*-clade *Wolbachia* are closely related to *w*Mel, globally polymorphic in *D. melanogaster*. We use draft *Wolbachia* and mitochondrial genomes to demonstrate that *D. yakuba*-clade phylogenies for *Wolbachia* and mitochondria tend to follow host nuclear phylogenies. However, roughly half of *D. santomea* individuals, sampled both inside and outside of the São Tomé hybrid zone, have introgressed *D. yakuba* mitochondria. Both mitochondria and *Wolbachia* possess far more recent common ancestors than the bulk of the host nuclear genomes, precluding cladogenic *Wolbachia* acquisition. General concordance of *Wolbachia* and mitochondrial phylogenies suggests that horizontal transmission is rare, but varying relative rates of molecular divergence complicate chronogram-based statistical tests. Loci that cause CI in *w*Mel are disrupted in *D. yakuba*-clade *Wolbachia*; but, a second set of loci predicted to cause CI are located in the same WO prophage region. These alternative CI loci seem to have been acquired horizontally from distantly related *Wolbachia*, with transfer mediated by flanking *Wolbachia*-specific ISWpi1 transposons.

## INTRODUCTION

Endosymbiotic *Wolbachia* bacteria infect many arthropods (Bouchon *et al*. 1998; Hilgenboecker *et al*. 2008), including about half of all insect species (Werren and Windsor 2000; Weinert *et al*. 2015). *Wolbachia* often manipulate host reproduction, facilitating spread to high frequencies within host species (Laven 1951; Yen and Barr 1971; Turelli and Hoffmann 1991; Rousset *et al*. 1992; O’Neill *et al*. 1998; Weeks *et al*. 2007; Kriesner *et al*. 2016; Turelli *et al*. 2018). In *Drosophila*, reproductive manipulations include cytoplasmic incompatibility (CI) and male killing (Hoffmann *et al*. 1986; Hoffmann and Turelli 1997; Hurst and Jiggins 2000). CI reduces the egg hatch of uninfected females mated with *Wolbachia*-infected males, and recent work has demonstrated that WO prophage-associated loci cause CI (Beckmann and Fallon 2013; Beckmann *et al*. 2017; LePage *et al*. 2017; Beckmann *et al*. 2019). Although reproductive manipulations are common, some *Wolbachia* show little or no reproductive manipulation (*e.g*., *w*Mel in *D. melanogaster*, Hoffmann 1988; Hoffmann *et al*. 1994; Kriesner *et al*. 2016; *w*Mau in *D. mauritiana*, Giordano *et al*. 1995; Meany *et al*. 2019; *w*Au in *D. simulans*, Hoffmann *et al*. 1996; *w*Suz in *D. suzukii* and *w*Spc in *D. subpulchrella*, Hamm *et al*. 2014; Cattel *et al*. 2018). These *Wolbachia* presumably spread by enhancing host fitness in various ways, with some support for viral protection, fecundity enhancement, and supplementation of host nutrition (Weeks *et al*. 2007; Teixeira *et al*. 2008; Hedges *et al*. 2008; Brownlie *et al*. 2009; Martinez *et al*. 2014; Gill *et al*. 2014; Moriyama *et al*. 2015; Kriesner and Hoffmann 2018). Better understanding of *Wolbachia* effects, transmission and evolution should facilitate using *Wolbachia* for biocontrol of human diseases by either transforming vector populations with virus-blocking *Wolbachia* (*e.g*., McMeniman *et al*. 2009; Hoffmann *et al*. 2011; Schmidt *et al*. 2017; Ritchie 2018) or using male-only releases of CI-causing *Wolbachia* to suppress vector populations (Laven 1967; O’Connor *et al*. 2012).

There is a burgeoning literature on *Wolbachia* frequencies and dynamics in natural populations (*e.g*., Kriesner *et al*. 2013; Kriesner *et al*. 2016; Cooper *et al*. 2017; Bakovic *et al*. 2018; Meany *et al*. 2019), but fewer studies elucidate the modes and time scales of *Wolbachia* acquisition by host species (O’Neill *et al*. 1992; Rousset and Solignac 1995; Huigens *et al*. 2004; Baldo *et al*. 2008; Raychoudhury *et al*. 2009; Ahmed *et al*. 2015; Schuler *et al*. 2016; Turelli *et al*. 2018). Sister hosts could acquire *Wolbachia* from their most recent ancestors. Such cladogenic acquisition seems to be the rule for the obligate *Wolbachia* found in filarial nematodes (Bandi *et al*. 1998); and there are also examples in at least two insect clades, *Nasonia* wasps (Raychoudhury *et al*. 2009) and *Nomada* bees (Gerth and Bleidorn 2016). In contrast, infections can be relatively young and acquired through introgressive or horizontal transfer (Raychoudhury *et al*. 2009; Schuler *et al*. 2016; Conner *et al*. 2017; Turelli *et al*. 2018).

Comparisons of host nuclear and mitochondrial genomes with the associated *Wolbachia* genomes enable discrimination among cladogenic, introgressive, and horizontal acquisition (Raychoudhury *et al*. 2009; Turelli *et al*. 2018; see Figure S1). Concordant nuclear, mitochondrial, and *Wolbachia* cladograms—including consistent divergence-time estimates for all three genomes—support cladogenic acquisition and co-divergence of host and *Wolbachia* lineages. Concordant *Wolbachia* and mitochondrial phylogenies and consistent *Wolbachia* and mitochondrial divergence-time estimates that are more recent than nuclear divergence support introgressive acquisition. In this case, mitochondrial and *Wolbachia* relationships may or may not recapitulate the host phylogeny. Finally, if *Wolbachia* diverged more recently than either nuclear or mitochondrial genomes, horizontal transfer (or paternal transmission; Hoffmann and Turelli 1988; Turelli and Hoffmann 1995) is indicated. This is often associated with discordance between host and *Wolbachia* phylogenies (O’Neill *et al*. 1992; Rousset and Solignac 1995; Turelli *et al*. 2018).

Introgressive *Wolbachia* acquisition may be common in *Drosophila*. About half of all closely related *Drosophila* species have overlapping geographical ranges and show pervasive evidence of reinforcement (Coyne and Orr 1989, 1997; Yukelevich 2012; Nosil 2013), indicating that hybridization must be common (Turelli *et al*. 2014). Several instances of sporadic contemporary hybridization and interspecific gene flow have been documented in the genus (Carson *et al*. 1989; Shoemaker *et al*. 1999; Jaenike *et al*. 2006; Kulathinal *et al*. 2009; Garrigan *et al*. 2012; Brand *et al*. 2013; Matute and Ayroles 2014; Lohse *et al*. 2015); but only two stable hybrid zones have been well described, and both involve *D. yakuba*-clade species. In west Africa, *D. yakuba* hybridizes with endemic *D. santomea* on the island of São Tomé, and with *D. teissieri* on the island of Bioko (Lachaise *et al*. 2000; Llopart *et al*. 2005; Comeault *et al*. 2016; Cooper *et al*. 2018). The ranges of *D. yakuba* and *D. teissieri* overlap throughout continental Africa, but contemporary hybridization has not been observed outside of Bioko (Cooper *et al*. 2018). Genomic analyses support both mitochondrial and nuclear introgression in the *D. yakuba* clade (Lachaise *et al*. 2000; Bachtrog *et al*. 2006; Llopart *et al*. 2014; Turissini and Matute 2017; Cooper *et al*. 2018), and the *Wolbachia* infecting all three species (*w*San, *w*Tei, and *w*Yak) are at intermediate frequencies and identical with respect to commonly used typing loci (Lachaise *et al*. 2000; Charlat *et al*. 2004; Cooper *et al*. 2017).

Horizontal *Wolbachia* transmission has been repeatedly demonstrated since its initial discovery by O’Neill *et al*. (1992) (*e.g*., Baldo *et al*. 2008; Schuler *et al*. 2016), and it may be common in some systems (*e.g*., Huigens *et al*. 2000; Huigens *et al*. 2004; Ahmed *et al*. 2015; Li *et al*. 2017). Phylogenomic analyses indicate recent horizontal *Wolbachia* transmission, on the order of 5,000–27,000 years, among relatively distantly related *Drosophila* species of the *D. melanogaster* species group (Turelli *et al*. 2018); but there is little evidence for non-maternal transmission within *Drosophila* species (Richardson *et al*. 2012; Turelli *et al*. 2018). Horizontal *Wolbachia* transfer could occur via a vector (Vavre *et al*. 2009; Ahmed *et al*. 2015) and/or via shared food sources during development (Huigens *et al*. 2000; Li *et al*. 2017). Rare paternal *Wolbachia* transmission has been documented in *D. simulans* (Hoffmann and Turelli 1988; Turelli and Hoffmann 1995). However, the most common mode of *Wolbachia* acquisition by *Drosophila* species (and most other hosts) remains unknown, and all three modes seem plausible in the *D. yakuba* clade (Lachaise *et al*. 2000; Bachtrog *et al*. 2006; Cooper *et al*. 2017).

Distinguishing acquisition via introgression versus horizontal or paternal transmission requires estimating phylogenies and relative divergence times of mtDNA and *Wolbachia* (Raychoudhury *et al*. 2009; Conner *et al*. 2017; Turelli *et al*. 2018). To convert sequence divergence to divergence-time estimates requires understanding relative rates of divergence for mtDNA, nuclear genomes and *Wolbachia*. If each genome followed a constant-rate molecular clock, taxa with cladogenic *Wolbachia* transmission, such as *Nasonia* wasps (Raychoudhury *et al*. 2009) and *Nomada* bees (Gerth and Bleidorn 2016), would provide reference rates for calibration. However, like Langley and Fitch (1974), Turelli *et al*. (2018) found significantly varying relative rates of *Wolbachia* and mtDNA divergence. This variation, which we document across *Drosophila*, confounds attempts to understand *Wolbachia* acquisition, as discussed below.

The discovery of weak CI in the *D. yakuba* clade (Cooper *et al*. 2017) motivates detailed comparative analysis of loci associated with CI (CI factors or *cifs*) in WO prophage regions of *Wolbachia* genomes. Beckmann and Fallon (2013) first associated wPip_0282 and wPip_0283 proteins in *w*Pip-infected *Culex pipiens* with *Wolbachia*-modified sperm. Later work confirmed that these proteins induce toxicity and produce rescue when expressed/co-expressed in *Saccharomyces cerevisiae* (Beckmann *et al*. 2017), lending support to a toxin-antidote model of CI (Beckmann *et al*. 2019; but see, Shropshire *et al*. 2019). Homologs of these genes in *w*Mel (WD0631 and WD0632) recapitulate CI when transgenically expressed in *D. melanogaster* (LePage *et al*. 2017), and transgenic expression of WD0631 in *D. melanogaster* rescues CI (Shropshire *et al*. 2018). A distantly related WO-prophage-associated pair, present in *w*Pip, *w*Pip_0294 and *w*Pip_0295, causes similar toxicity/rescue in *S. cerevisiae* (Beckmann *et al*. 2017), and causes CI when placed transgenically into a *D. melanogaster* background (unpublished data presented by Mark Hochstrasser at the 2018 *Wolbachia* meeting in Salem, MA). We adopt Beckmann *et al*. (2019)’s nomenclature, which assigns names based on enzymatic activity of the predicted toxin [deubiquitylase (DUB) and nuclease (Nuc)], with superscripts denoting focal *Wolbachia* strains when needed. Specifically, we refer to *w*Pip_0282-*w*Pip_0283 and the wPip_0294-wPip_0295 pairs as *cidA-cidB^w^*^Pip^ and *cinA-cinB^w^*^Pip^, respectively; and we refer to WD0631-WD632 as *cidA-cidB^w^*^Mel^. This distinguishes *cid* (CI-inducing DUB) from *cin* (CI-inducing Nuc) pairs, with the predicted antidote and toxin denoted “A” and “B”, respectively (Beckmann *et al*. 2019). We acknowledge ongoing disagreement in the literature and direct readers to Beckmann *et al*. (2019) and Shropshire *et al*. (2019) for details. However, none of these debates on terminology or mechanism affect our findings.

Here, we use host and *Wolbachia* genomes from the *D. yakuba* clade to demonstrate introgressive and horizontal *Wolbachia* acquisition. General concordance of mitochondrial and *Wolbachia* phylogenies indicates that horizontal acquisition is rare within this clade. However, tests involving divergence-time estimates are complicated by varying relative rates and patterns of *Wolbachia*, mtDNA, and nuclear sequence divergence, as illustrated by data from more distantly related *Drosophila* (Clark *et al*. 2007). Finally, we demonstrate that *cid* loci underlying CI in closely related *w*Mel (LePage *et al*. 2017) are disrupted in all *D. yakuba*-clade *Wolbachia*. However, these *Wolbachia* also contain a set of *cin* loci, absent in *w*Mel, but very similar to those found in *w*Pip (Beckmann *et al*. 2017), a B-group *Wolbachia* strain that diverged 6–46 million years ago (mya) from A-group *w*Yak and *w*Mel (Werren *et al*. 1995; Meany *et al*. 2019). This is the first discovery of two sets of loci implicated in CI co-occuring within the same prophage region. Several analyses implicate *Wolbachia*-specific insertion sequence (IS) transposable elements, specifically ISWpi1, in the horizontal transfer of these loci (and surrounding regions) between distantly related *Wolbachia*. Horizontal movement of incompatibility factors between prophage regions of *Wolbachia* variants adds another layer to what is already known about horizontal movement of prophages within and between *Wolbachia* variants that themselves move horizontally between host species.

## MATERIALS AND METHODS

### Genomic data

The *D. yakuba*-clade isofemale lines included in our study were sampled over several years in west Africa (Comeault *et al*. 2016; Turissini and Matute 2017; Cooper *et al*. 2017). Each line within each species used in our analyses exhibits little nuclear introgression (< 1%) (Turissini and Matute 2017), and no hybrids were included. Reads from *D. yakuba* (*N* = 56), *D. santomea* (*N* = 11), and *D. teissieri* (*N* = 13) isofemale lines were obtained from the data archives of Turissini and Matute (2017) and aligned to the *D. yakuba* nuclear and mitochondrial reference genomes (Clark *et al*. 2007) with bwa 0.7.12 (Li and Durbin 2009), requiring alignment quality scores of at least 50. Because many of the read archives were single end, all alignments were completed using single-end mode for consistency.

#### mtDNA

Consensus mtDNA sequences for each of the 80 lines were extracted with samtools v. 1.3.1 and bcftools v 1.3.1 (Li 2011). Coding sequences for the 13 protein-coding genes were extracted, based on their positions in the *D. yakuba* reference. We also extracted the 13 protein-coding genes from additional unique *D. yakuba* (*N* = 28) and *D. santomea* (*N* = 15) mitochondrial genomes (Llopart *et al*. 2014), and from the *D. melanogaster* reference (Hoskins *et al*. 2015). Genes were aligned using MAFFT v. 7 and concatenated (Katoh and Standley 2013). Lines identical across all 13 genes were represented by one sequence for the phylogenetic analyses.

#### Wolbachia

Reads from each of the 80 Turissini and Matute (2017) lines were aligned using bwa 0.7.12 (Li and Durban 2009) to the *D. yakuba* reference genome (Clark *et al*. 2007) combined with the *w*Mel reference genome (Wu *et al*. 2004). We calculated the average depth of coverage across the *w*Mel genome. We considered lines with < 1× coverage uninfected and lines with >10× coverage infected. No lines had between 1× and 10× coverage. To test our genomic analyses of infection status, we used a polymerase chain reaction (PCR) assay on a subset of lines. We extracted DNA using a “squish” buffer protocol (Gloor *et al*. 1993), and infection status was determined using primers for the *Wolbachia*-specific *wsp* gene (Braig *et al*. 1998; Baldo *et al*. 2006). We amplified a *Drosophila*-specific region of chromosome *2L* as a positive control (Kern *et al*. 2015) (primers are listed in Supplemental Material, Table S1). For each run, we also included a known *Wolbachia*-positive line and a water blank as controls.

To produce a draft *Wolbachia* “pseudoreference” genome for each species, we first determined the isofemale line from each species that produced the greatest average coverage depth over *w*Mel. We trimmed the reads with Sickle v. 1.33 (Joshi and Fass 2011) and assembled with ABySS v. 2.0.2 (Jackman *et al*. 2017). *K* values of 51, 61…91 were tried. Scaffolds with best nucleotide BLAST matches to known *Wolbachia* sequences with E-values less than 10^−10^ were extracted as the draft *Wolbachia* assembly. For each species, the assembly with the highest N50 and fewest scaffolds was kept as our *Wolbachia* pseudoreference genome for that host species, denoted *w*Yak, *w*San, and *w*Tei (Table S2). To assess the quality of these three draft assemblies, we used BUSCO v. 3.0.0 to search for homologs of the near-universal, single-copy genes in the BUSCO proteobacteria database (Simão *et al*. 2015). For comparison, we followed Conner *et al*. (2017) and performed the same search on the reference genomes for *w*Mel (Wu *et al*. 2004), *w*Ri (Klasson *et al*. 2009), *w*Au (Sutton *et al*. 2014), *w*Ha and *w*No (Ellegaard *et al*. 2013) (Table S3).

Using these draft *Wolbachia* pseudoreference genomes, reads from all other genotypes were aligned to the *D. yakuba* reference (nuclear and mitochondrial) plus the species-specific *Wolbachia* draft assembly with bwa 0.7.12 (Li and Durbin 2009). Consensus sequences were extracted with samtools v. 1.3.1 and bcftools v. 1.3.1 (Li 2011).

To test whether the choice of *Wolbachia* pseudoreference influenced the *Wolbachia* draft sequence obtained for each infected line, we arbitrarily selected three infected lines of each host species and aligned reads from each of those lines independently to our *w*Yak, *w*San, and *w*Tei pseudoreferences. This resulted in 27 alignments (9 per host species). Among the three *Wolbachia* consensus sequences generated for each line, we assessed single-nucleotide variants between different mappings of the same line within loci used for downstream analyses. Irrespective of the pseudoreference used, we obtained identical *Wolbachia* sequences. This indicates the robustness of our approach to generating population samples of draft *Wolbachia* genomes, without having a high-quality *Wolbachia* reference genome from any of the host species. Because Llopart *et al*. (2014) released only assembled mitochondrial genomes, we could not assess the *Wolbachia* infection status of their lines.

### Loci for phylogenetic and comparative-rate analyses

#### Wolbachia genes

For each phylogenetic analysis of *Wolbachia* data, the draft genomes were annotated with Prokka v. 1.11 (Seemann 2014), which identifies homologs to known bacterial genes. To avoid pseudogenes and paralogs, we used only genes present in a single copy and with identical lengths in all of the sequences analyzed. Genes were identified as single copy if they uniquely matched a bacterial reference gene identified by Prokka v. 1.11. By requiring all homologs to have identical length in all of our draft *Wolbachia* genomes, we removed all loci with indels across any of our sequences.

Given that many loci accumulate indels over time, the number of loci included in our phylogenetic analyses depended on the number of strains included. We first estimated phylograms for the A-group *Wolbachia* from: the *D. yakuba*-clade, *w*Mel from *D. melanogaster* (Wu *et al*. 2004), *w*Inc from *Drosophila incompta* (Wallau *et al*. 2016), *w*Suz from *D. suzukii* (Siozios et al. 2013), *w*Ana from *D. anannasae* (Choi *et al*. 2015), *Wolbachia* that infect *Nomada* bees (*w*NFe, *w*NPa, *w*NLeu, and *w*NFa; Gerth and Bleidorn 2016), and *Wolbachia* that infect *D. simulans* (*w*Ri, *w*Au and *w*Ha; Klasson *et al*. 2009; Sutton *et al*. 2014; Ellegaard *et al*. 2013). We also analyzed distantly related B-group *Wolbachia*: *w*No from *D. simulans* (Ellegaard *et al*. 2013), *w*Pip_Pel from *Culex pipiens* (Klasson *et al*. 2008), and *w*AlbB from *Aedes albopictus* (Mavingui *et al*. 2012). To increase our phylogenetic resolution within the *D. yakuba Wolbachia* clade (by increasing the number of loci), we estimated a phylogram that included only *D. yakuba*-clade *Wolbachia* and *w*Mel. Our phylogram with both A- and B-group *Wolbachia* included 146 genes (containing 115,686 bp), and our phylogram that included only *D. yakuba*-clade *Wolbachia* and *w*Mel included 643 genes (containing 644,586 bp).

We also constructed an absolute chronogram to estimate divergence between *D. yakuba*-clade *Wolbachia* strains. To illustrate how divergence-time estimates vary with changing patterns of molecular divergence, we estimated two additional *Wolbachia* chronograms: one that considered only the *w*Mel reference plus the two most-diverged *w*Mel lines included in Richardson *et al*. (2012), and one that included these three *w*Mel variants and our *D. yakuba*-clade *Wolbachia*. Our chronogram that included only *D. yakuba*-clade *Wolbachia* included 678 genes (containing 695,118 bp), our chronogram that included only *w*Mel *Wolbachia* included 692 genes (containing 709,599 bp), and our chronograms with *w*Mel plus *D. yakuba*-clade *Wolbachia* included 621 genes (containing 624,438 bp). Independent estimates were obtained using relaxed-clock analyses with Γ(2,2) and Γ(7,7) branch-rate priors. We also estimated divergence between *D. yakuba*-clade *Wolbachia* and *w*Mel using a strict-clock analysis, corresponding to Γ(*n*,*n*) as *n* → ∞.

#### mtDNA and nuclear genes from diverse *Drosophila*

To assess variation in the relative rates of divergence of mitochondrial and nuclear genes, we analyzed the canonical 12 *Drosophila* genomes (Clark *et al*. 2007) plus *D. suzukii* (Chiu *et al*. 2013). We excluded *D. sechellia* and *D. persimilis* which show evidence of introgression with *D. simulans* and *D. pseudoobscura*, respectively (Kulathinal *et al*. 2009; Schrider *et al*. 2018; Brand *et al*. 2013). Coding sequences for 20 nuclear genes used in the analyses of Turelli *et al*. (2018) (*aconitase, aldolase, bicoid, ebony, enolase, esc, g6pdh, glyp, glys, ninaE, pepck, pgi, pgm, pic, ptc, tpi, transaldolase, white, wingless*, and *yellow*) were obtained from FlyBase for each species. The genes were aligned with MAFFT v. 7 (Katoh and Standley 2013). Coding sequences for the 13 protein-coding mitochondrial genes in the inbred reference strains were also obtained from FlyBase for each species and were aligned with MAFFT v. 7 and concatenated.

### Phylogenetic analyses

All of our analyses used RevBayes v. 1.0.9 (Hohna *et al*. 2016), following the procedures of Turelli *et al*. (2018). For completeness, we summarize those methods below. For additional details on the priors and their justifications, consult Turelli *et al*. (2018). Four independent runs were performed for each phylogenetic tree we estimated; and in all cases, the runs converged to the same topologies. Nodes with posterior probability less than 0.95 were collapsed into polytomies.

#### *Wolbachia* phylograms

We estimated a phylogram for A- and B-group *Wolbachia* and for only *D. yakuba*-clade and *w*Mel *Wolbachia* using the same methodology as Turelli *et al*. (2018). We used a GTR + Γ model with four rate categories, partitioning by codon position. Each partition had an independent rate multiplier with prior Γ(1,1) (i.e., Exp(1)), as well as stationary frequencies and exchangeability rates drawn from flat, symmetrical Dirichlet distributions (i.e., Dirichlet(1,1,1…). The model used a uniform prior over all possible topologies. Branch lengths were drawn from a flat, symmetrical Dirichlet distribution, thus they summed to 1. Since the expected number of substitutions along a branch equals the branch length times the rate multiplier, the expected number of substitutions across the entire tree for a partition is equal to the partition’s rate multiplier.

#### *Wolbachia* chronograms

We first created a relaxed-clock relative chronogram with the root age fixed to 1 using the GTR + Γ model, partitioned by codon position, using the same birth-death prior as Turelli *et al*. (2018). Each partition had an independent rate multiplier with prior Gamma(1,1), as well as stationary frequencies and exchangeability rates drawn from flat, symmetrical Dirichlet distributions. The branch-rate prior for each branch was Γ(2,2), normalized to a mean of 1 across all branches (Table S4). We also tried a strict-clock tree and a relaxed-clock tree with branch-rate prior Γ(7,7), which produced no significant differences. We used the scaled distribution Γ(7,7) × 6.87 × 10^−9^ to model substitutions per third-position site per year. This transforms the relative chronogram into an absolute chronogram. This scaled distribution was chosen to replicate the upper and lower credible intervals of the posterior distribution estimated by Richardson *et al*. (2012), assuming 10 *Drosophila* generations per year, normalized by their median substitution-rate estimate. Branch lengths in absolute time were calculated as the relative-branch length times the third-position rate multiplier divided by the substitutions per third-position site per year estimate above.

We illustrate how divergence-time estimates depend on changing patterns of *Wolbachia* molecular evolution observed over different time scales, specifically the relative rates of third-site substitutions versus first- and second-site substitutions. We compare divergence times estimated for variants within *melanogaster*-group host species, with a time scale of only hundreds or thousands of years, with the same divergence times estimated when more distantly related *Wolbachia*, with divergence times over tens of thousands of years, are included in the analyses. We present three separate analyses: one using only *D. yakuba*-clade *Wolbachia* variants, a second using only *w*Mel variants analyzed by Richardson *et al*. (2012), and a third simultaneously analyzing both sets of data.

#### *Drosophila* phylogeny for 11 references species

We estimated a phylogram from our 20 nuclear loci using the GTR + Γ model, with partitioning by gene and codon position. The model was identical to the *Wolbachia* phylogram model above except for the partitioning (there are too few *Wolbachia* substitutions to justify partitioning by gene).

#### mtDNA phylogeny for the 11 reference species

We estimated a phylogram from the 13 protein-coding mitochondrial loci using the GTR + Γ model, with partitioning only by codon position. The model was identical to the *Wolbachia* phylogram model above.

#### mtDNA chronogram for the *D. yakuba* clade

We estimated a relative chronogram for the *D. yakuba*-clade mitochondria from the 13 protein-coding loci using the GTR + Γ model, partitioning only by codon position. To test the sensitivity of our results to priors, we ran a strict clock, a relaxed clock with a Γ(7,7) branch-rate prior, and a relaxed clock with a Γ(2,2) branch-rate prior, corresponding to increasing levels of substitution-rate variation across branches. The model was identical to the *Wolbachia* chronogram model above, except that we did not transform it into an absolute chronogram.

#### Ratios of divergence rates for mtDNA versus nuclear loci

To quantify ratios of mtDNA to nuclear substitution rates, we estimated relative substitution rates for host nuclear genes versus mtDNA using the GTR + Γ model. The (unrooted) topology was fixed to the consensus topology for the nuclear and mitochondrial data. The data from 20 nuclear loci were partitioned by locus and codon position, the mtDNA data were partitioned only by codon position. All partitions shared the same topology, but the nuclear partitions were allowed to have branch lengths different from the mtDNA partitions. The sum of the branch lengths for each partition was scaled to 1. Assuming concurrent nuclear-mtDNA divergence (because we used only species showing no evidence of introgression), we imposed the same absolute ages for all nodes of the nuclear and mtDNA chronograms. For each nuclear locus, the third-position rate-ratio for the mtDNA versus nuclear genomes was calculated as: (mitochondrial branch length × mitochondrial 3^rd^ position rate multiplier)/(nuclear branch length × nuclear 3^rd^ position rate multiplier). We summarized the relative rates of mtDNA versus nuclear substitutions along each branch using the arithmetic average of the 20 ratios obtained from the individual nuclear loci.

#### Introgressive versus horizontal *Wolbachia* transfer–concordance of phylograms

Horizontal transfer of *Wolbachia* was initially detected by discordance between *Wolbachia* and host phylogenies (O’Neill *et al*. 1992). When considering samples within species and closely related species, a comparison of mitochondrial and *Wolbachia* phylogenies provides a natural test for horizontal transmission. We first look for significant discrepancies between strongly supported nodes in the mitochondrial versus *Wolbachia* phylogenies. To test for less obvious differences, we follow Richardson *et al*. (2012) and compute Bayes Factors to assess the support for models that assume that mitochondrial versus *Wolbachia* follow the same topology versus distinct phylogenies. For these calculations, we omit san-Quija37 due to its clear discordance.

We calculated the marginal likelihood in RevBayes for two models; one with a shared mitochondrial and *Wolbachia* phylogeny, another with independent topologies. The mtDNA and *Wolbachia* were each partitioned by codon position, for a total of six partitions. In the shared phylogeny model, all six share the same topology; in the independent model, mitochondria and *Wolbachia* have separate topologies. All priors were the same as the *Wolbachia* phylogram model. Lines that were identical across the mtDNA and *Wolbachia* were collapsed into one sample. We ran two independent replicates of each model with 50 stepping stones per run. The Bayes factor is computed as the difference between the marginal likelihoods of each model.

#### Introgressive versus horizontal *Wolbachia* transfer–relative rates

Within species, the phylogenies of mitochondria and *Wolbachia* are fully resolved. Thus we consider an alternative approach for distinguishing between introgressive versus horizontal transfer that depends on estimating divergence times for *Wolbachia* genomes and host mtDNA. This is complicated by variation in the relative rates of mtDNA versus *Wolbachia* divergence and by systematic changes over time in the relative rates of substitutions at the three codon positions for *Wolbachia*. Following the procedures in Turelli *et al*. (2018), we estimated the mtDNA-to-*Wolbachia* third-position substitution-rate ratio for each branch in the *D. yakuba* clade. For each analysis, we computed the marginal likelihood of the model where all branches shared the same ratio and the same model except allowing different ratios on each branch. We then calculated Bayes factors (i.e. differences in the log of the marginal likelihoods) to determine which model was favored. We repeated these analyses after including three *w*Mel-infected *D. melanogaster* lines with the *D. yakuba*-clade dataset. Because *D. yakuba*-clade species and *D. melanogaster* do not produce fertile hybrids (Sánchez and Santamaria 1997; Turissini *et al*. 2017), introgressive transfer of *Wolbachia* is not possible between these species. Hence, including *w*Mel provides a control for our proposed test of introgressive transfer. For these controls, we calculated the mitochondrial versus *Wolbachia* substitution-rate ratio for the interspecific branches separately, regardless of whether the shared rate-ratio model was favored.

### *Wolbachia* loci associated with CI

*Wolbachia* that infect all three *D. yakuba*-clade hosts cause weak intra-and interspecific CI (Cooper *et al*. 2017), but the genetic basis of CI in this clade remains unknown. We used tBLASTN to search for *cif* homologs in each of our *D. yakuba*-clade *Wolbachia* assemblies, querying *cidA-cidB* variants and the *cinA-cinB^w^*^Ri^ pair (wRi_006720 and wRi_006710) found in *w*Ri and some *w*Ri-like *Wolbachia* (Beckmann and Fallon 2013; Turelli *et al*. 2018), and the *cinA-cinB^w^*^Pip^ pair (Beckmann and Fallon 2013; Beckmann *et al*. 2017; LePage *et al*. 2017; Lindsey *et al*. 2018). We identified both *cid* and *cin* homologs, but we found no close matches to the *cinA-cinB^w^*^Ri^ pair (denoted Type II *cif* loci by Lepage *et al*. 2017) in any of our genomes. For all of the samples, we extracted consensus sequences from our assemblies and alignments for the *cidA-cidB^w^*^Yak-clade^ and *cinA-cinB^w^*^Yak-clade^ gene pairs. The genes were aligned with MAFFT v. 7 (Katoh and Standley 2013). We examined variation in these regions relative to *cidA-cidB^w^*^Mel^ and *cinA-cinB^w^*^Pip^ respectively.

Because we unexpectedly identified homologs of *cinA-cinB* in all *w*Yak-clade *Wolbachia*, we took additional measures to understand the origin and placement of these loci in the genomes. The *cinA-cinB^w^*^Yak-clade^ open reading frames are located on a ∼11,500 bp scaffold in the fragmented *w*Yak assembly (*w*Yak scaffold “702380” from ABySS output––we use quotes to indicate names assigned by ABySS). The first ∼4,000 bp of this scaffold contain *cinA* and *cinB* genes that have ∼97% identity with those in *w*Pip. We performed a BLAST search using all contigs in the *w*Yak assembly as queries against the *w*Mel genome (Camacho *et al*. 2009). The *cinA-cinB^w^*^Yak-clade^ scaffold was placed on the *w*Mel genome (∼617,000–623,000) adjacent to and downstream of the *w*Yak contig containing *cidA-cidB^w^*^Yak-clade^ loci (*w*Yak contig “187383”). However, only ∼7,000 bp of the 11,500 bp of the scaffold containing *cinA*-*cinB^w^*^Yak-clade^ align to this region of *w*Mel. The ∼4,000 bp sequence that contains the ORFs for *cinA*-*cinB^w^*^Yak-clade^ has no significant hit against the *w*Mel genome.

To verify the placement of the scaffold containing *cinA*-*cinB^w^*^Yak-clade^, we first performed targeted assembly using the *cinA*-*cinB^w^*^Yak-clade^ contig and the adjacent contigs as references (Hunter *et al*. 2015). Iterative targeted assembly often extends assembled scaffolds (Langmead and Salzberg 2012). Reads are mapped to the target of interest, and the mapped reads are then assembled using SPADES (Bankevich *et al*. 2012; Nurk *et al*. 2013). The newly assembled scaffolds serve as the target in a subsequent round of mapping. The procedure is repeated until no new reads are recruited to the assembled scaffold from the previous round. The extended scaffolds enabled us to merge flanking *w*Yak contigs downstream of the *cinA*-*cinB^w^*^Yak-clade^ scaffold but failed to connect the fragment with the *cidA*-*cidB^w^*^Yak-clade^ contig. We used PCR to amplify the intervening region (Supplemental Information, Table S1), and Sanger sequencing to evaluate this product (Sanger *et al*. 1977).

Subsequent mapping of paired-end reads to the merged scaffolds confirmed the correct order and orientation of the contigs containing the *cidA-cidB^w^*^Yak-clade^ and *cinA-cinB^w^*^Yak-clade^ genes. Observed, uncorrected, pairwise distances between focal regions in *w*Yak, *w*Mel, and *w*Pip and between focal regions in *w*Yak, *w*Pip, *w*AlbB, and *w*Nleu were calculated using a sliding window (window size = 200 bp, step size = 25 bp).

### Data Availability

The *Wolbachia* assemblies (BioProject PRJNA543889, Accession VCEF00000000) and Sanger sequences (MK950151 and MK950152) are archived in GenBank. All three assemblies were manually corrected using Sanger sequence to accurately represent variation in the *cin* region. All relevant code will be uploaded to Dryad prior to publication.

## RESULTS

### Unidirectional introgression of *D. yakuba* mitochondria into *D. santomea*

Previous analyses suggest that *D. yakuba* and *D. santomea* carry very similar mtDNA due to ongoing hybridization and introgression (Lachaise *et al*. 2000; Bachtrog *et al*. 2006; Llopart *et al*. 2014). However, our mtDNA relative chronogram (Figure 1), based on mitochondrial whole-proteome data sampled from throughout the ranges of all three species, supports three mtDNA clades that largely agree with the nuclear topology—*D. teissieri* mtDNA sequences are outgroup to sister *D. yakuba* and *D. santomea* mtDNA (Figure 1). Consistent with introgression, 12 out of 26 *D. santomea* isofemale lines have *D. yakuba*-like mtDNA (indicated by blue branches and letters in Figure 1). Yet, our samples showed no evidence of *D. santomea* mtDNA introgressed into *D. yakuba* and no mtDNA introgression involving *D. teissieri*. The 12 *D. santomea* isofemales with introgressed *D. yakuba*-like mtDNA were sampled from both within (*N* = 8) and outside (*N* = 4) the well-described Pico de São Tomé hybrid zone (Llopart et al. 2014; Turissini and Matute 2017). The *D. santomea* hybrid-zone samples are indicated by “HZ” in Figure 1 in blue, and they are found throughout the clade that includes all *D. yakuba* mtDNA. This suggests that hybridization occurs in other areas of the island or that introgressed *D. santomea* genotypes migrate outside of the hybrid zone.

**Figure 1.**
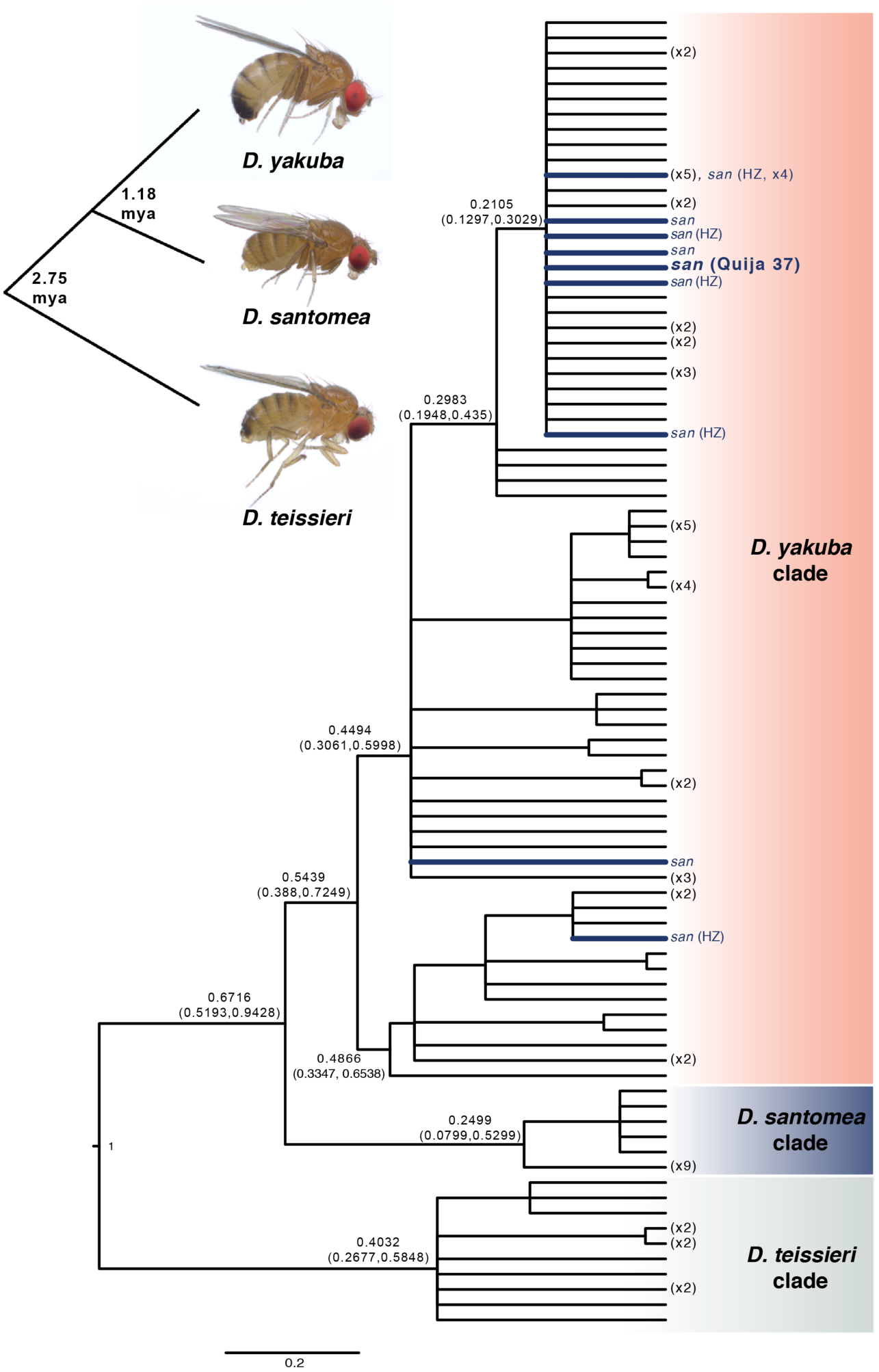
Bayesian relative chronogram for the *D. yakuba* clade mtDNA. The root age was fixed to 1. Nodes with posterior probability less than 0.95 were collapsed into polytomies. (The numbers at the tips show the number of genotypes collapsed.) Node labels are the point estimates and 95% credible intervals for the relative ages. Tips and branches showing *D. santomea* individuals with *D. yakuba* mtDNA are colored blue, those sampled from the Pico de São Tomé hybrid zone are denoted by HZ. *D. santomea* line Quija 37 showing evidence of horizontal or paternal *Wolbachia* transmission is in bold blue. The chronogram was estimated from the thirteen protein-coding mitochondrial genes. A representation of host nuclear relationships with photos of host females at the tips is inset, with the divergence times estimated by Turissini and Matute (2017) superimposed.

Previous analysis of nuclear genomes (Turissini and Matute 2017) also found more introgression from *D. yakuba* into *D. santomea* when populations of *D. yakuba* from near the Gulf of Guinea, Cameroon, and Kenya were included; however, excluding Cameroon and Kenya indicated similar amounts of introgression in each direction. Matings between *D. santomea* females and *D. yakuba* males are rare relative to the reciprocal cross (Coyne *et al*. 2002, Matute 2010). When these matings do occur, F1 females produce fewer progeny than do F1 females produced by *D. yakuba* females and *D. santomea* males (Matute and Coyne 2010). F1 hybrids produced by *D. santomea* females also have a shortened lifespan (Matute and Coyne 2010), due to copulatory wounds inflicted by *D. yakuba* males during mating (Kamimura 2012). These observations are consistent with our finding of preferential introgression of *D. yakuba* mitochondria into *D. santomea* backgrounds.

### *Wolbachia* frequencies and draft genomes

As expected, we found *Wolbachia* in all three *D. yakuba*-clade species sampled by Turissini and Matute (2017). For *D. yakuba*, 21 of 56 lines were infected, yielding an infection frequency of *p* = 0.36, with 95% binomial confidence interval (0.25, 0.51). For *D. santomea*, 10 of 11 were infected, *p* = 0.91 (0.59, 1.0); and for *D. teissieri*, 11 of 13 were infected, *p* = 0.85 (0.55, 0.98). Additional frequency estimates are reported in Cooper *et al*. (2017), which found that *Wolbachia* frequencies vary through time and space in west Africa. All PCR tests for *Wolbachia* matched our coverage-based genomic analyses of infection status.

The average coverage across the genome, calculated as the total number of bases aligned to the *w*Mel reference divided by its length, was 1,940 for our *w*Yak pseudoreference genome (yak-CY17C), 40 for *w*San (san-Quija630.39), and 489 for *w*Tei (teis-cascade_4_2). These pseudoreference genomes were included in the phylogram that includes A- and B-group *Wolbachia* (Figure 2), and they cluster with the *Wolbachia* from their respective hosts (Figure 3), as expected if introgression is rare. In general, the *w*San data are lower quality due to the relatively small size of the *D. santomea* libraries (Turissini and Matute 2017). The scaffold count, N50, and total assembly sizes are reported in Table S2.

**Figure 2.**
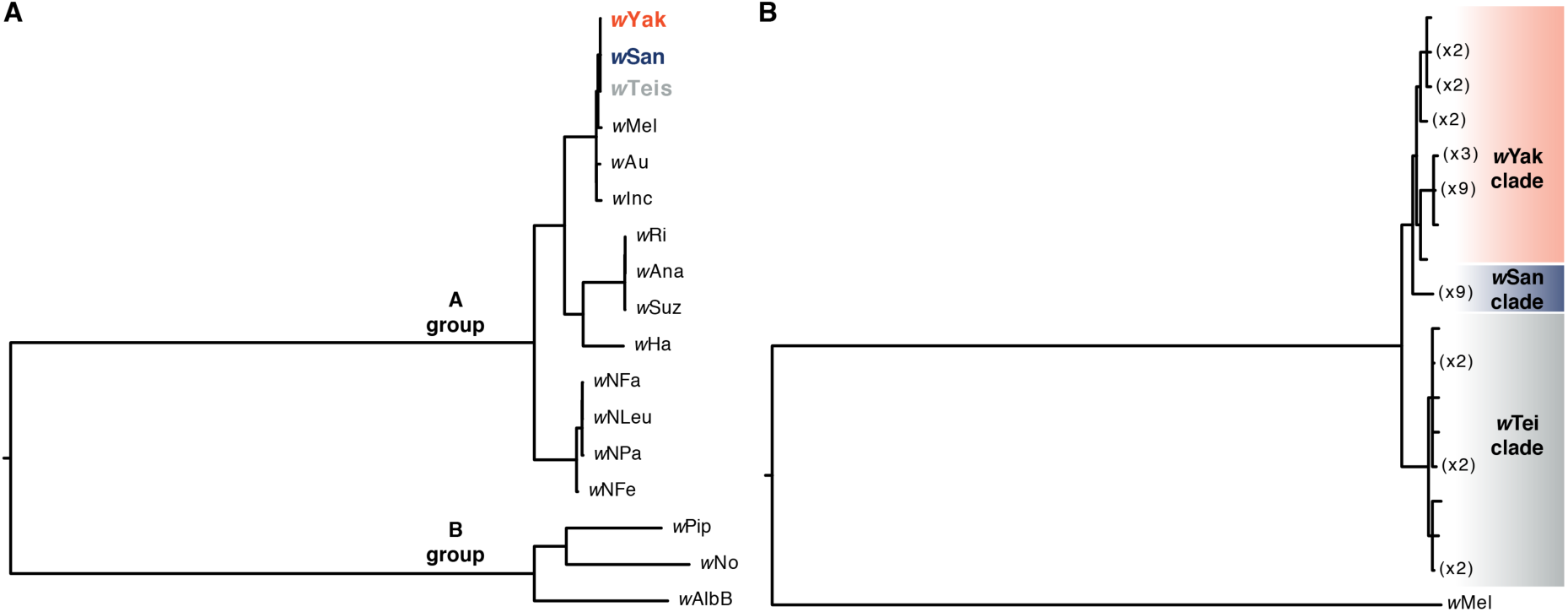
A) Bayesian phylogram placing the *D. yakuba*-clade *Wolbachia* in the context of other A-, and also B-group, *Wolbachia*. *w*Pip, *w*No, and *w*AlbB belong to *Wolbachia* group B, while *D. yakuba* clade-*Wolbachia*, *w*Mel, *w*Au, *w*Inc, *w*Ri, *w*Ana, *w*Suz, *w*Ha, and *Nomada Wolbachia* belong to group A. The phylogram was estimated with 146 single-copy genes of identical length in each genome, spanning 115,686 bp. The limited number of genes meeting our criteria precluded resolving relationships of *D. yakuba*-clade *Wolbachia*. B) Bayesian phylogram of *w*Yak, *w*San, *w*Tei, and outgroup *w*Mel. The phylogram was estimated with 643 single-copy genes of identical length in each genome, spanning 644,586 bp. These more extensive data provide increased phylogenetic resolution of the *D. yakuba*-clade *Wolbachia*. For both phylograms, nodes with posterior probability less than 0.95 were collapsed into polytomies.

**Figure 3.**
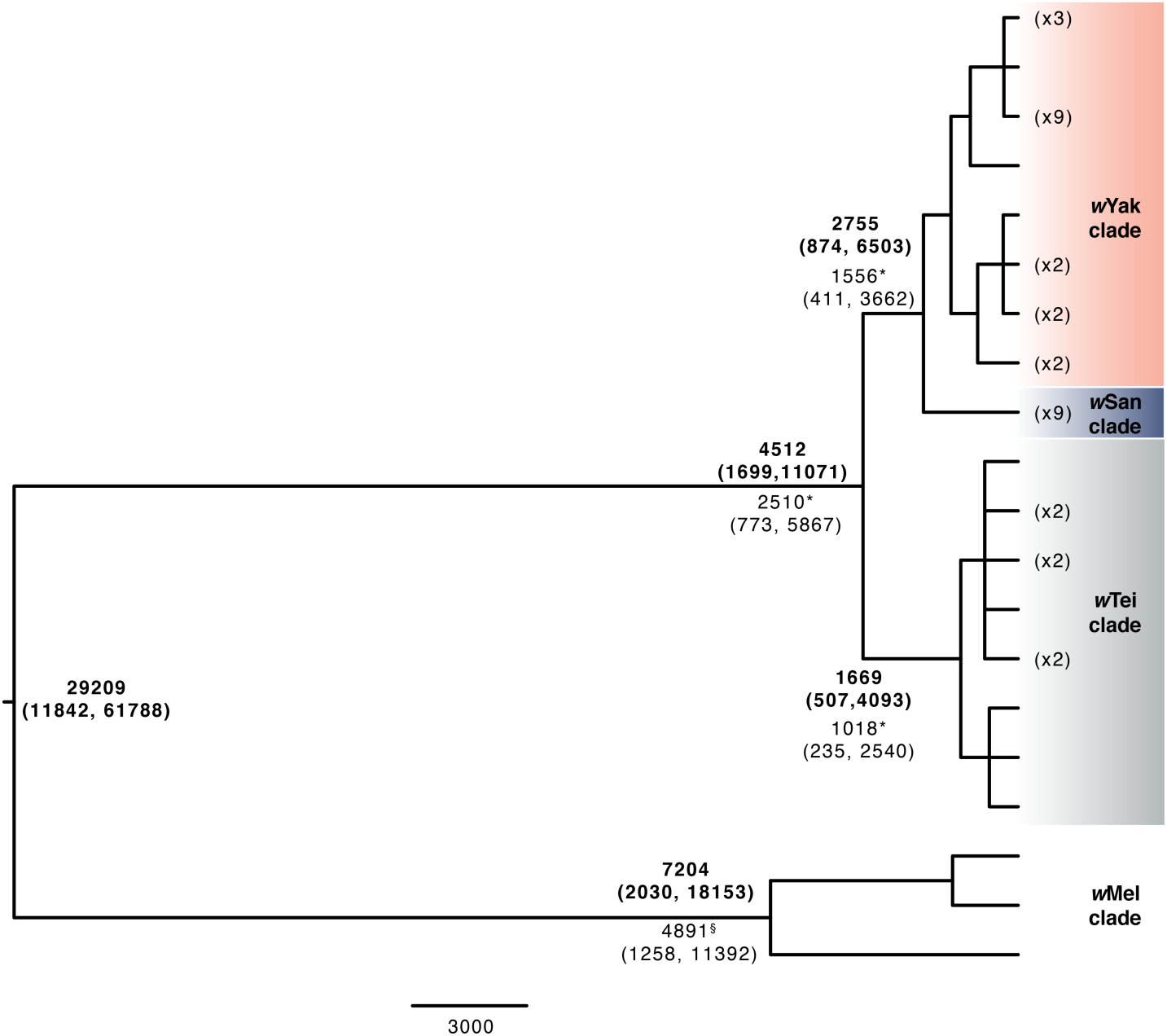
Bayesian chronogram for the *D. yakuba*-clade *Wolbachia* plus three *w*Mel genomes with absolute age estimates based on the calibration in Richardson *et al*. (2012) in bold. The chronogram was estimated with 621 single-copy genes of identical length in all of the genomes, spanning 624,438 bp. Nodes with posterior probability less than 0.95 were collapsed into polytomies. Node labels are the point estimates and 95% credible intervals for the ages. We also constructed two other chronograms: one that included only *D. yakuba*-clade *Wolbachia* and one that included only *w*Mel *Wolbachia*. The divergence estimates for the chronogram including only *D. yakuba*-clade *Wolbachia* (*) and only *w*Mel *Wolbachia* (^§^) are superimposed to illustrate that the inclusion of both groups in the chronogram alters the absolute divergence estimates within each group, although credible intervals overlap in each case. Our chronogram with only *D. yakuba*-clade *Wolbachia* included 678 genes (containing 695,118 bp), and our chronogram including only *w*Mel *Wolbachia* included 692 genes (709,599 bp).

### Introgressive and horizontal *Wolbachia* transfer among hosts

Our phylogram that includes A- and B-group *Wolbachia* places the three weak-CI-causing *D. yakuba*-clade *Wolbachia* together and sister to *w*Mel in the A group (Figure 2A). In contrast, *D. simulans* carries diverse *Wolbachia* that do (*w*Ri, *w*Ha, *w*No) and do not (*w*Au) cause CI, spanning *Wolbachia* groups A (*w*Ri, *w*Ha, *w*Au) and B (*w*No). All *w*San, *w*Yak, and *w*Tei *Wolbachia* variants included in our analysis are identical at the MLST loci often used to type *Wolbachia* variants (Cooper *et al*. 2017; Baldo *et al*. 2006). Due to the relatively small number of genes that meet our inclusion criteria (146 genes across 115,686 bp), the resulting phylogram does not resolve relationships among *w*San, *w*Yak, and *w*Tei (Figure 2A). Our phylogram that includes only *D. yakuba*-clade and *w*Mel *Wolbachia* (based on 643 genes across 644,586 bp) resolves these relationships, placing *w*Tei variants outgroup to sister *w*San and *w*Yak in distinct clades (Figure 2B). We observe 0.0039% third-position pairwise differences between *w*Tei and *w*Yak and between *w*Tei and *w*San, and 0.0017% between *w*San and *w*Yak. The third-position pairwise differences between any of the *D. yakuba*-clade *Wolbachia* and *w*Mel is 0.11%. For reference, the third-position pairwise difference between *w*Yak and *w*Ri is 2.9%, whereas *w*Ri and *w*Suz differ by only 0.014% (Turelli *et al*. 2018).

Our absolute chronogram that includes *w*Mel and *D. yakuba*-clade *Wolbachia* also indicates three monophyletic groups (Figure 3), whose topology––(*w*Tei, (*w*Yak, *w*San))––agrees with the nuclear topology of the hosts (Turissini and Matute 2017). However, one *D. santomea* line, *D. santomea* Quija 37, sampled from the south side of São Tomé (Matute 2010), carries *Wolbachia* identical to the other eight *w*San samples across 695,118 bp, yet its mtDNA is nested in the *D*. *yakuba* mtDNA cluster (bold blue in Figure 1). This single example of *Wolbachia*-mtDNA phylogenetic discordance can be explained by either horizontal *Wolbachia* transfer or paternal transfer of *Wolbachia* or mtDNA. While rare, paternal *Wolbachia* transmission has been observed in other *Drosophila* (Hoffmann and Turelli 1988; Turelli and Hoffmann 1995), as has paternal transmission of mtDNA (Kondo *et al*. 1990). The only other *Wolbachia*-screened *D. santomea* isofemale line with introgressed mtDNA was *Wolbachia* uninfected.

In Figure 3, we present alternative estimates of the divergence times for *Wolbachia* within and between *D. yakuba*-clade species, within *D. melanogaster*, and between the *D. yakuba* clade and *D. melanogaster*. The estimates from the chronogram that includes all of the indicated *Wolbachia* variants are in bold. To demonstrate how these time estimates depend on the sequences included in the analyses (through estimates of relative divergence for each codon position), estimates that included only *D. yakuba*-clade *Wolbachia* or only *w*Mel variants are superimposed and not in bold. These estimates were all obtained using a relaxed-clock analysis with a Γ(2,2) branch-rate prior (results using Γ(7,7) and a strict clock are mentioned below). When considering only the *D. yakuba*-clade *Wolbachia* (Figure 3), we estimate the root at 2,510 years (95% credible interval = 773 to 5,867 years) and the *w*Yak-*w*San split at 1,556 years (95% credible interval = 411 to 3,662 years). The *w*Mel chronogram estimates the most recent common ancestor (MRCA) of *w*Mel at 4,890 years ago (95% credible interval = 1,258 to 11,392 years). Our *w*Mel result is consistent with the Richardson *et al*. (2012) estimate of 8,008 years (95% credible interval = 3,263, 13,998). The alternative branch-rate Γ(7,7) prior has little effect on these divergence-times estimates, with credible intervals that overlap with those produced using a Γ(2,2) branch-rate prior (Figure 3 and Table S5). In contrast, Figure 3 shows that including more divergent variants influences our time estimates, although the credible intervals are overlapping in each case. Table S4 shows how estimates of the relative amounts of the divergence for the three codon positions vary across the data sets. The underlying model for divergence times assumes constant relative rates at the three positions. Hence, changes in relative rates affect divergence-time estimates.

As noted above, our estimates of the time of the MRCA for *Wolbachia* within the *D. yakuba* clade or within *D. melanogaster* are relatively insensitive to whether the branch-rate prior is Γ(2,2) or Γ(7,7). The choice of genes used to estimate divergence also has little effect—maximal distance between *D. yakuba*-clade strains was identical for analyses that included only the *D. yakuba*-clade (*N* = 678 genes) or *w*Yak-clade plus *w*Mel (*N* = 621 genes) gene sets (*w*Yak-*w*Tei distance = 0.0035%). The estimated divergence time between *w*Mel and the *D. yakuba*-clade *Wolbachia* is less robust, although again the credible intervals generated using alternative branch-rate priors are overlapping (Figure 3 and Table S5). A strict-clock analysis produces an estimate of 72,612, with 95% credible interval (23,412, 132,276), again overlapping with our relaxed-clock estimates. An alternative point estimate can be obtained from pairwise differences. Averaging over the *D. yakuba*-clade sequences, we observe an average third-position difference of 0.107% between the *D. yakuba*-clade *Wolbachia* and *w*Mel, or 0.0535% from tip to root. The “short-term evolutionary rate” of divergence within *w*Mel estimated by Richardson *et al*. (2012) produces a point estimate of 78,000 years, which overlaps with the credible interval of our relaxed-tree estimate with a Γ(7,7) branch-rate prior (Table S5). This molecular-clock estimate closely agrees with our strict-clock chronogram analysis (Figure S2). Given that these estimates are much shorter than the divergence time between their reproductively isolated hosts, horizontal *Wolbachia* transmission must have occurred. As discussed below, the horizontal transmission probably involved intermediate hosts.

#### mtDNA and nuclear concordance across *Drosophila*

The topologies of the nuclear and mtDNA trees for the 11 reference *Drosophila* species are completely concordant (Figure 4) and agree with the neighbor-joining result presented in Clark *et al*. (2007). We find the average ratio of mtDNA to nuclear substitution rates across branches vary from about 1× to 10×, consistent with the results of Havird and Sloan (2016) across Diptera.

**Figure 4.**
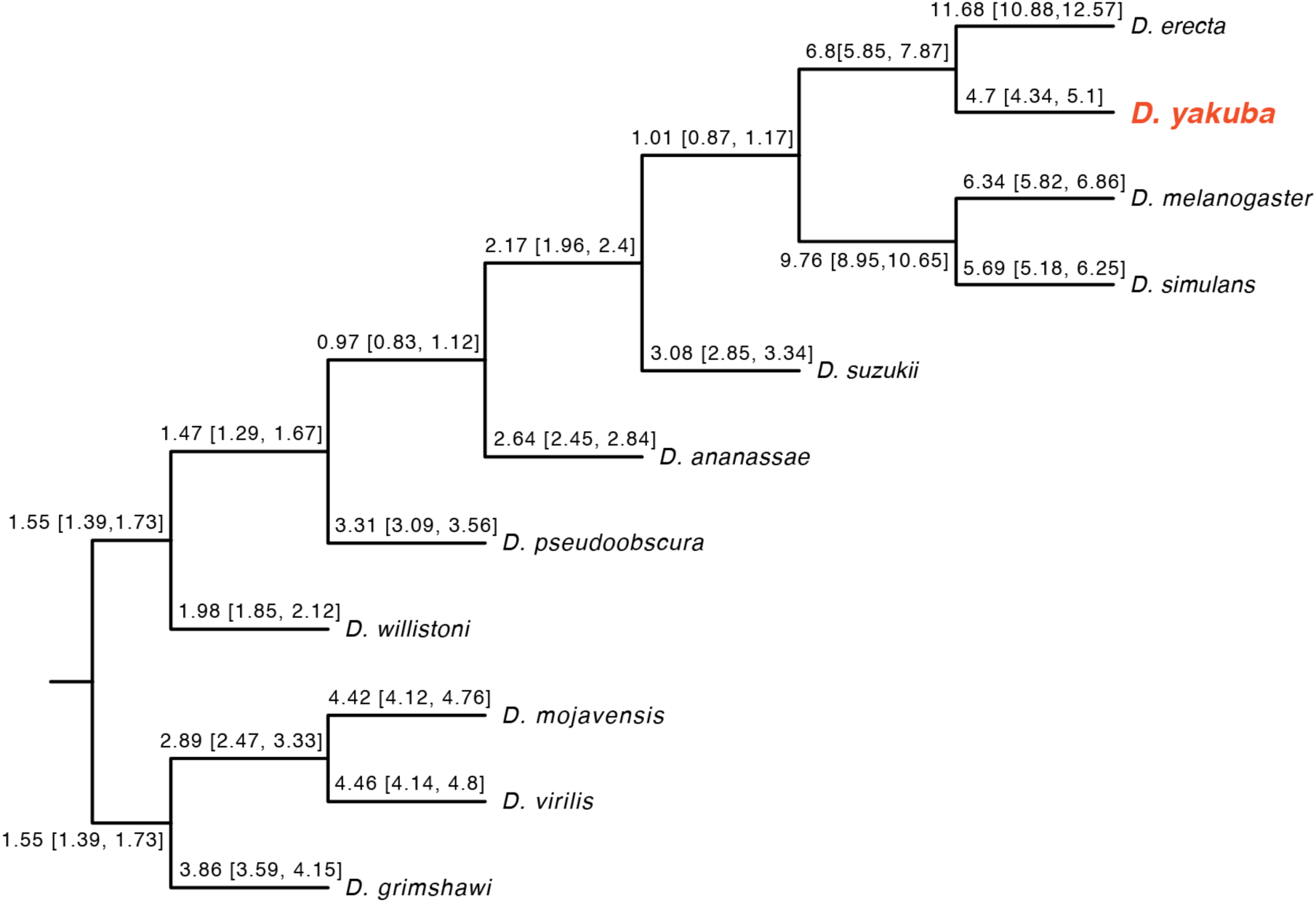
A Bayesian phylogram, with branch lengths fixed to 1, displaying relative rate variation for mtDNA versus nuclear loci. Branches are labeled with quartiles of the estimated third position mtDNA to third position nuclear substitution rate. The nuclear substitution rate was taken as the arithmetic mean third-position rate over the 20 loci used. The mitochondrial and nuclear phylograms are topologically identical when run separately.

#### mtDNA and *Wolbachia* concordance in the *D. yakuba* clade

The discordance between the placement of the mitochondria of *D. santomea* line Quija 37, which has mitochondria belonging to the clade associated *D. yakuba* and its *Wolbachia*, which is clearly *w*San, demonstrates horizontal or paternal transmission. Apart from this, we find no nodes in either the mitochondrial or *Wolbachia* tree with posterior probability at least 0.95 that is inconsistent with a strongly supported node (i.e., posterior probability at least 0.95) in the other tree (Figure S3). For a more refined test of concordance, we removed the Quija 37 line from our mitochondrial and *Wolbachia* data and computed Bayes Factors. The shared topology model was favored by a Bayes factor of e^55^, indicating strong support for concordance of the mitochondrial and *Wolbachia* phylogenies.

#### Variation in the ratio of mtDNA to *Wolbachia* substitution rates

Following the methods of Turelli *et al*. (2018) to look for different mitochondrial versus *Wolbachia* branch lengths, we also found no evidence of variation in the ratio of mtDNA:*Wolbachia* substitution rates within the *D. yakuba* clade (median 3^rd^ position ratio = 223, quartiles = 186 and 268); the model with all branches sharing the same ratio is favored over the model where each branch has its own ratio by a Bayes factor of e^7^ (Figure S4). For comparison, Turelli *et al*. (2018) found a median ratio of 566 within *D. suzukii* and 406 within *w*Ri-infected *D. ananassae*-subgroup species. These differences in relative rates are broadly compatible with the tenfold variance in relative rates noted above for nuclear versus mtDNA divergence across *Drosophila* species. Hence, our data seem compatible with *Wolbachia* transmission via introgression within the *D. yakuba* clade.

Although this result seems to strongly support purely maternal transmission of *Wolbachia* within species and introgressive transfer between species (with more recent introgression between the sister species *D. yakuba* and *D. santomea*), this interpretation is severely weakened by our “control” analysis that includes *w*Mel-infected *D. melanogaster* (for which introgression is impossible). Including the *D. melanogaster* data, the constant-ratio model was still favored over variable ratios by a Bayes factor of e^54^ (median 3^rd^ position ratio = 297, quartiles = 276 and 323). We discuss the implications of this anomalous result below. The key observation is that significantly declining rates of substitution for *Wolbachia* (and mtDNA, see Ho *et al*. 2005) over time, together with heterogeneity of relative rates (as illustrated in Figure 4), limit the power of relative substitution ratios to differentiate between introgression and horizontal transmission.

### Transposon-mediated transfer of CI factors independent of WO phage

In contrast to *w*Mel, which contains only the *cidA-cidB* gene pair (LePage et al. 2017), the *D. yakuba*-clade *Wolbachia*, which also belong to *Wolbachia* group A, have both *cidA-cidB* and a *cinA-cinB* pair homologous to CI loci originally identified in group-B *w*Pip (Beckmann *et al*. 2017). The A and B *Wolbachia* groups diverged about 6-46 mya (Werren *et al*. 1995; Meany *et al*. 2019). Among our *w*Yak-clade sequences, there are no single-nucleotide variants within any of the *cif* loci. The *cidB^w^*^Yak-clade^ locus has an inversion from amino acids 37–103 relative to the same region in *cidB^w^*^Mel^ in every *w*Yak-clade *Wolbachia* variant we analyzed. This introduces several stop codons, which might render this gene nonfunctional. On the other hand, RNA polymerase should still transcribe a complete polycistronic transcript. Therefore translation of an N-terminally truncated CidB*^w^*^Yak-clade^ protein cannot be ruled out. With the exception of a 236 bp tandem duplication in *cinB^w^*^Yak-clade^ (Figures 5 and 6), the sequence differences between *cinA^w^*^Yak-clade^ and *cinB^w^*^Yak-clade^ regions compared to *cinA^w^*^Pip^ and *cinB^w^*^Pip^ homologs are 1.35% and 0.59%, respectively. In contrast, the average difference between *w*Yak and *w*Pip genomes across all 194 genes (161,655 bp) present in single copy and with identical lengths in both genomes is 11.60%. Conversely, outside of the prophage regions, *w*Mel and *w*Yak differ by about 1%. This is consistent with data that indicate WO phage regions have a different evolutionary history than the bulk of the *Wolbachia* genome (LePage *et al*. 2017; Lindsey *et al*. 2018). The 236 bp tandem duplication in the *cinB^w^*^Yak-clade^ introduces a frame shift in the transcript at position 588. It is unclear whether the *cinB^w^*^Yak-clade^ protein retains functionality.

**Figure 5.**
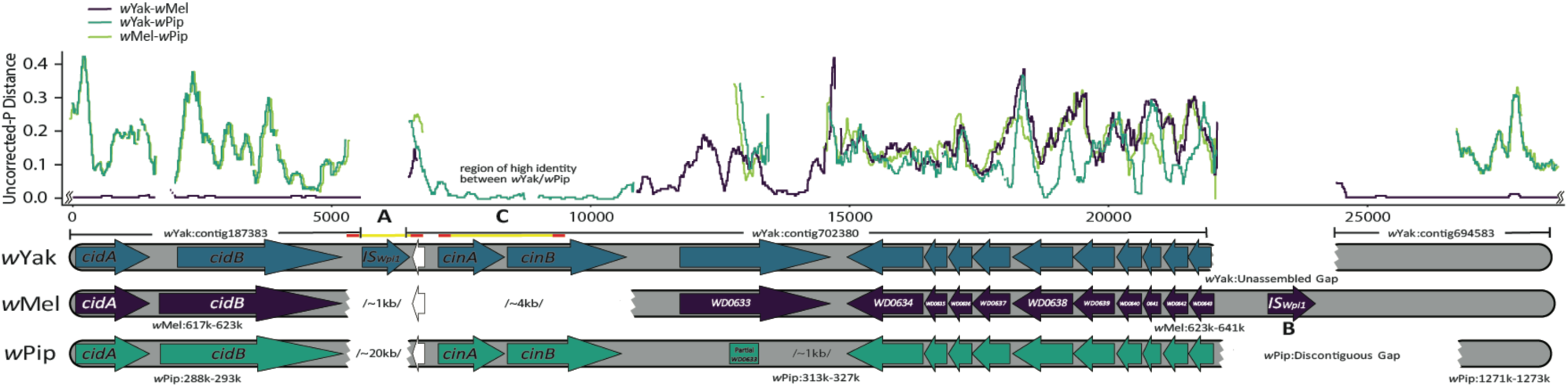
The gene structure and observed, uncorrected, pairwise distances among *w*Yak, *w*Mel, and *w*Pip *Wolbachia* spanning the region of the WO prophage island where the hypothesized transfer occurred between the ISWpi1 element “A” present in *w*Yak and the ISWpi1 sequence labeled “B” in *w*Mel. Over the region shown, *w*Yak, *w*San and *w*Tei are identical. The difference between *w*Yak and *w*Mel is less than 1% between the contigs (*w*Yak “187383” and *w*Yak “694583”) flanking the region containing the *cinA*-*cinB* homologs. These homologs assembled on the first ∼4,000 bp of the contig *w*Yak “702380”, but they do not appear to exist in *w*Mel. A BLAST search against the NCBI nucleotide collection using contig *w*Yak “702380” as a query indicated the *cinA*-*cinB* homologs share high identity with group-B *w*Pip *Wolbachia* (∼97%). The remainder of the *w*Yak contig is approximately equally divergent from both *w*Mel and *w*Pip, consistent with the entire region being transferred from an unknown *Wolbachia* variant that had acquired the *cinA-cinB* homologs. Under this scenario, ISWpi1 elements mediate the transfer to *w*Yak via excision of elements “A” and “B” along with the intervening DNA from the donor and subsequent homologous recombination between IS sequences “A” and “B” and WO phage island genes in *w*Yak. The ISWpi1 element “A” present in *w*Yak, but not in either *w*Mel or *w*Pip, was spanned using Sanger sequencing with primers anchored on unique sequence in the flanking contigs (Sanger sequences are denoted by yellow lines with primer locations denoted by red flanking regions, Table S1). Additionally, the presence of a 236 bp tandem duplication in *cinB* (“C”) was confirmed using Sanger sequencing. The ISWpi1 sequence labeled “B” is assembled in *w*Mel and corresponds to *w*Mel#9 (Cordaux 2008) but could not be assembled or spanned with Sanger sequencing in the *w*Yak genome. White gene models (arrows) indicate a partial transposase gene present in the WO prophage island, truncated in *w*Yak. Pairwise differences were calculated using a sliding window (window size = 200 bp, step size = 25 bp).

**Figure 6.**
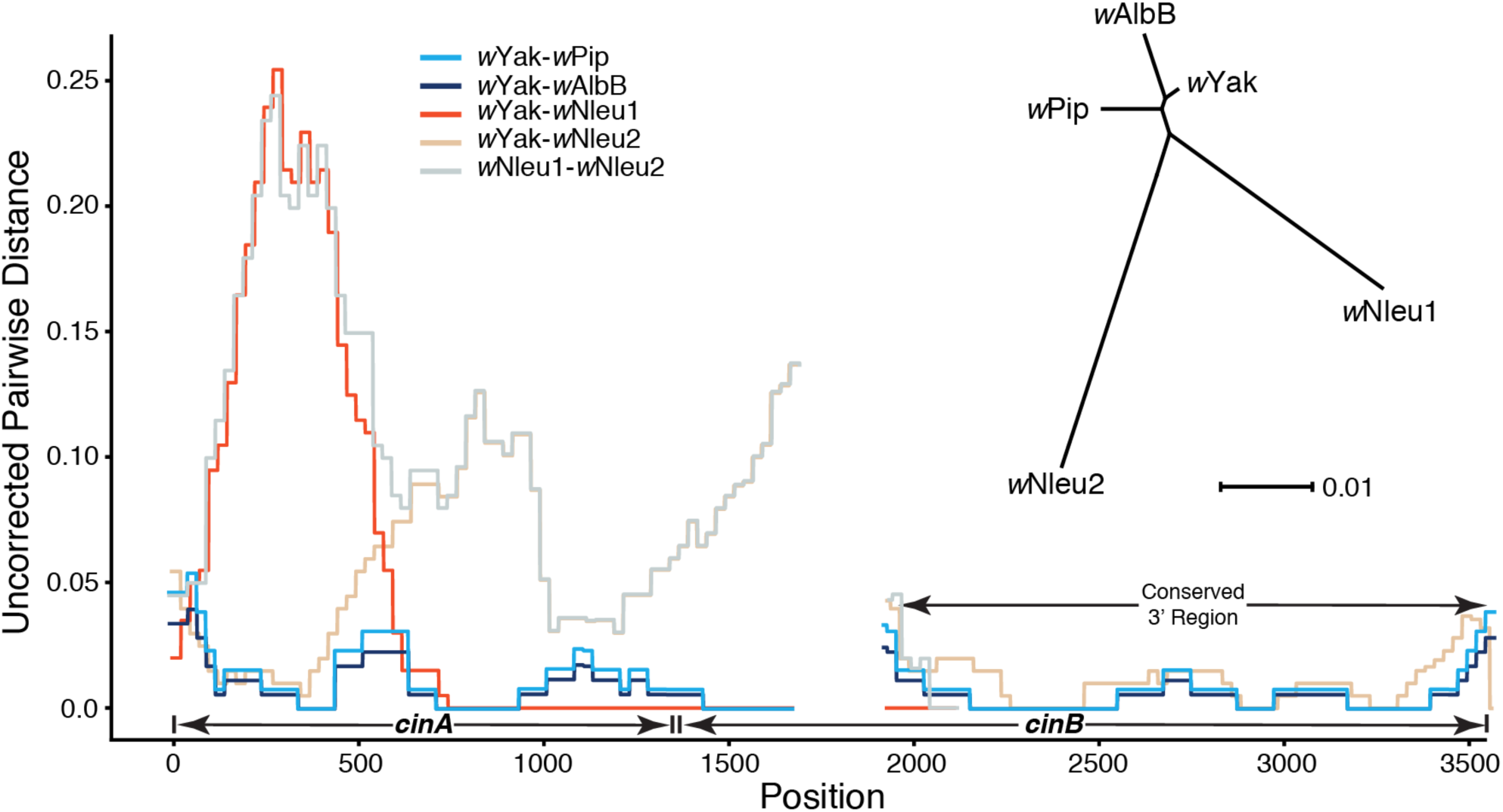
The observed pairwise differences between *cinA* and *cinB* copies in *w*Yak-clade *Wolbachia* and those in *w*AlbB and *w*Pip, not including flanking regions, are shown. We also plot the pairwise differences between the two copies of these loci found in *Nomada Wolbachia* (denoted *w*Nleu1 and *w*Nleu2); we chose *w*Nleu to represent the *Nomada*-clade *Wolbachia* because this strain contains the longest assembled contigs for both *cinA-cinB* copies. Homologous *cinA-cinB* copies have relatively low divergence (inset unrooted tree), with the highest divergence in the 5’ end of this region. The most highly diverged region among any of the copies is the first ∼750 bp of *w*Nleu1, with pairwise differences between *w*Nleu1 and *w*Yak reaching ∼25% in some windows, with similar divergence between *w*Nleu1 and *w*Nleu2 in this same region. Pairwise differences abruptly change 750 bp from the start of the *cinA* gene with the remaining ∼1,150 bp of the assembled region having a single difference from the *w*Yak sequence (only ∼1,900 bp assembled into a single contig in the *w*Nleu1 copy). This pattern suggests a recent transfer/recombination event from the same unknown donor of the *w*Yak *cinA-cinB* copy. The gap at position 1705-1940 represents a tandem duplication present in *w*Yak. The unrooted tree was generated using RAxML 8.2.9 (Stamatakis 2014), and the representative *cinA-cinB* gene copies were obtained using BLAST to search all *Wolbachia* genomes in the NCBI assemblies database (https://www.ncbi.nlm.nih.gov/assembly/?term=Wolbachia).

The absence of *cin* loci in *w*Mel, combined with the similarity of this region between *w*Yak and distantly related *w*Pip, led us to assess how these *Wolbachia* acquired *cin* loci. Targeted assembly extended the scaffold *w*Yak “702380” containing the *cin* loci (Figure 5) but did not definitively place it relative to the *w*Mel genome. BLAST searches indicated that contig *w*Yak “187383” was likely flanked by *w*Yak “702380”. PCR primers designed in both contigs amplified the intervening region (labeled “A” in Figure 5), confirming we have discovered the first WO prophage with two sets of *cif* loci. Subsequent Sanger sequencing revealed that this region contains an ISWpi1 element (Supplemental Information), found in many *Wolbachia* genomes, but not present at this location in the *w*Mel reference sequence (Cordeaux *et al*. 2008).

IS elements encode a transposase gene that mediates their movement (Chandler and Mahillon 2002). ISWpi1 elements are related to the IS5 family, which seems to be restricted to *Wolbachia* (Duron *et al*. 2005; Cordaux *et al*. 2008). ISWpi1 elements occur in more than half of the *Wolbachia* strains that have been evaluated, including *w*Yak and *w*Mel (Cordaux *et al*. 2008); and these elements occur at variable copy number, potentially facilitating horizontal transmission of ISWpi1 elements and their intervening sequence between *Wolbachia* variants/strains (Cordaux *et al*. 2008). Following placement of contig *w*Yak “702380” relative to *w*Mel, we aligned these regions and calculated pairwise differences along the chromosome using a sliding window.

In addition to containing the *cinA-cinB* loci, absent in *w*Mel, contig *w*Yak “702380” is on average about 10% different from *w*Mel. In contrast, most of *w*Yak is less than 1% different from *w*Mel (for instance, across the 650,559 bp used in our phylogenetic analyses, *w*Mel and *w*Yak differ by only 0.09%). Downstream of the *cinA-cinB^w^*^Yak-clade^ region, targeted assembly enabled us to join several more contigs. The junctions were corroborated by mapping paired-end reads (Langmead and Salzberg 2012) and visually inspecting the resulting bam files around joined contigs for reads spanning the new junctions and for concordant read-pair mappings. However, our attempts to fully bridge the gap downstream of the *cinA-cinB^w^*^Yak-clade^ genes via targeted assembly, scaffolding, and PCR were unsuccessful (see “*w*Yak unassembled gap” in Figure 5). The unassembled region in *w*Yak contains an ISWpi1 element in *w*Mel, Mel #9 (labeled “B” in Figure 5; Cordaux 2008). Although not part of our *w*Yak assembly, homologs of this ISWpi1 element appear in assemblies of this region of the WO prophage in several other A-group *Wolbachia* (Cordaux *et al*. 2008; Bordenstein and Bordenstein 2016), specifically *w*Inc from *D. incompta* (Wallau *et al*. 2016) and *w*Ri from *D. simulans* (see Figure 2). In both, we find orthologs to Mel#9 ISWpi1 with corresponding flanking sequence assembled, indicating that this IS element probably occurs in this unassembled region of *w*Yak, *w*San, and *w*Tei. The *w*Yak sequence between these two ISWpi1 elements is highly diverged from *w*Mel in comparison to the rest of the *w*Yak genome (about 10% difference versus 1%). It therefore seems plausible that this region experienced a horizontal transfer event in the ancestor of *w*Yak, *w*Tei, and *w*San, mediated by the flanking ISWpi1 elements.

We conjectured that horizontal transfer occurred via the excision of the two ISWpi1 elements and the intervening DNA from the donor followed by homologous recombination within the IS elements. To assess the plausibility of this scenario, we used BLAST to compare *cinA-cinB^w^*^Pip^ genes against all published *Wolbachia* genomes in the NCBI assembly database (https://www.ncbi.nlm.nih.gov/assembly/?term=Wolbachia). We found homologs of the *cinA-cinB^w^*^Pip^ in the group-A *Wolbachia* associated with *Nomada* bees (Gerth and Bleidorn 2016). An unrooted tree of the ORFs for *cinA* and *cinB* (Figure 6) indicates that these genes in *w*Yak, *w*San, and *w*Tei are more similar to *cinA-cinB* from group-B *w*Pip and *w*AlbB than to *cinA-cinB* found in the fellow group-A *Wolbachia* associated with *Nomada* bees (*w*NFe, *w*NPa, *w*NLeu, and *w*NFa), which each harbor two different *cinA-cinB* copies. This indicates that *cinA-cinB^w^*^Yak-clade^ were acquired via a horizontal transfer event across *Wolbachia* groups B and A that is independent of the event(s) that placed *cinA-cinB* in the *Wolbachia* associated with *Nomada* bees, suggesting repeated transfers of *cin* loci.

The two *cinA-cinB* copies (denoted *w*NLeu1 and *w*NLeu2 in Figure 6) in *Nomada Wolbachia* are nearly as distinct from each other as they are from the homologs in *w*Pip, *w*AlbB, and *w*Yak (∼7% diverged from these strains, Figure 6). However, among the four *Wolbachia*-infected *Nomada* species, the orthologs are very similar, with *cinA^w^*^NLeu1^ having only 0–0.15% pairwise differences among the four strains and *cinA^w^*^NLeu2^ having 0–0.56% pairwise differences. (Reconstructions for *cinB* gene copies were more complicated as the *cinB^w^*^NLeu1^ copy fails to assemble into a single contig on the 3-prime end.) This pattern suggests that *w*NLeu1 and *w*NLeu2 *cin* copies were acquired by the common ancestor of the four *Nomada Wolbachia* strains analyzed, followed by cladogenic transfer across *Nomada* species (Gerth and Bleidorn 2016). The highly fragmented assemblies of the four *Nomada Wolbachia* strains, with duplicate copies confounding assembly, make it difficult to determine the relative positions of the *cinA* and *cinB* copies and if they are likewise flanked by ISWpi1 elements.

To determine whether these genes were potentially moved by ISWpi1 elements into the *Wolbachia* of the *D. yakuba* clade and *Nomada*, we searched each genome using both the *cinA-cinB^w^*^Yak-clade^ contig and the flanking ISWpi1 elements. Long repeated elements like ISWpi1 (916bp) break most short-read assemblies. Despite this, there is often a small fragment of the element, the length of the short read, on either end of the broken contig, indicative of the repeat element being responsible for terminating contig extension. We looked for the footprint of these elements at the edges of the contigs on which the *cin* genes were found. We found ISWpi1 elements in the region flanking both *cinA^w^*^NLeu1^ and *cinA^w^*^NLeu2^ copies, consistent with our *w*Yak assembly in which we verified the ISWpi1 element with Sanger sequencing. These data support a role for ISWpi1 in the acquisition of the *cinA-cinB* genes by the *Wolbachia* in the *D. yakuba* clade and the *Nomada* bees. We conjecture that future work will fully confirm ISWpi1 in the horizontal movement of incompatibility loci between *Wolbachia*.

## DISCUSSION

Our results indicate introgressive and horizontal *Wolbachia* acquisition in the *D. yakuba* clade. Evidence for horizontal *Wolbachia* transfer here and elsewhere (Turelli *et al*. 2018) suggests that double infections must be common, even if ephemeral. Such double infections provide an opportunity for ISWpi1 transposable elements to mediate horizontal transfer of incompatibility loci among divergent *Wolbachia*. Importantly, our results highlight that incompatibility factors may move independently of prophage, as evidenced by our discovery of the first prophage documented to have two sets of *cif* loci. We discuss these conclusions below.

### mtDNA Introgression

Our relative mitochondrial chronogram provides strong support for three mitochondrial clades, including a monophyletic *D. teissieri* clade that is outgroup to two sister clades: one clade consisting of mitochondria from 14 *D. santomea* individuals, and the other contains all *D. yakuba* mitochondria plus mitochondria from 12 *D. santomea* individuals. Our results suggest less mitochondrial introgression in the *D. yakuba* clade than a past report that used sequence data from only two mitochondrial loci (*COII* and *ND5*, 1777 bp) and reported a clade with mitochondria sampled from each species represented (Figure 3 in Bachtrog *et al*. 2006). (Using sequence data from only *COII* and *ND5*, we can replicate this result, indicating data from additional loci are needed to add resolution.) Our results also agree with recent work that demonstrated little nuclear introgression in the *D. yakuba* clade (Turissini and Matute 2017).

Our results broadly agree with the results produced by Llopart *et al*. (2014) who assessed mitochondrial introgression between *D. santomea* and *D. yakuba* using whole mitochondrial genomes. They generated a neighbor-joining tree that produced a clade consisting of all *D. yakuba* individuals and 10 *D. santomea* individuals, and this clade is sister to a clade with 6 *D. santomea* individuals; one *D. santomea* haplotype is outgroup to all other haplotypes included in their analysis (Figure 3 in Llopart *et al*. 2014). Nested within the mixed *D. yakuba* clade, Llopart *et al*. (2014) identified a “hybrid zone clade” that includes *D. yakuba* individuals sampled from São Tomé and from continental Africa and *D. santomea* individuals sampled from both within and outside the Pico de São Tomé hybrid zone (HZ). The sister *D. santomea* clade also contains both HZ and non-HZ individuals. Thus, their analysis and ours provide support for hybridization within and outside of the HZ, leading us to question both the existence of a HZ clade and the claim that these species “share the same mitochondrial genome” (Llopart *et al*. 2014). Instead, both our results and theirs suggest unidirectional introgression of *D. yakuba* mitochondria into *D. santomea*; we find 59% of *D. santomea* individuals having *D. yakuba*-like mitochondria and they found 46%.

Llopart *et al*. (2014) used a strict molecular clock to estimate the mitochondrial MRCA of *D. santomea* and *D. yakuba* at 10,792–17,888 years by calibrating their tree with the *D. yakuba-D. erecta* split, estimated at 10.4 mya (Tamura *et al*. 2004). A high level of mitochondrial saturation over time, with an expected value of 1.44 substitutions per synonymous site for the *D. yakuba-D. erecta* split, could influence this estimate (Llopart *et al*. 2014). Moreover, Ho *et al*. (2005) demonstrated that the mtDNA substitution rate resembles an exponential curve, with high short-term substitution rates that approach the mutation rate, then slowing to a long-term rate after about 1–2 million years of divergence, far younger than the inferred *D. yakuba-D. erecta* nuclear and mtDNA divergence. Hence, using the slow long-term *D. yakuba-D. erecta* calibration is likely to underestimate the more rapid rates of divergence experienced by *D. yakuba*-clade mtDNA, inflating divergence-time estimates (Ho *et al*. 2005). If we assume that *Wolbachia* and mitochondria were transferred by introgression, which our analyses support, our estimates in Figure 3 suggest that *D. santomea* and *D. yakuba* mtDNA diverged more recently, with our point estimates ranging from about 1,500 to 2,800 years.

### *Wolbachia* placement, divergence, and acquisition

#### Wolbachia placement

Despite their similarity, *w*San, *w*Tei, and *w*Yak form monophyletic clades with *w*Tei outgroup to sisters *w*San and *w*Yak, recapitulating host relationships (Figure 1; Turissini and Matute 2017). *w*Mel is sister to the *D. yakuba*-clade strains in the A group (Figure 2), which also includes *D. simulans* strains (*w*Ri, *w*Au, and *w*Ha), *w*Ana, *w*Suz, and the *Nomada Wolbachia* (*w*NFa, *w*NLeu, *w*NPa, and *w*NFe). Our phylogram (Figure 2A) places *w*AlbB outgroup to *w*No and *w*Pip strains that diverged from A-group *Wolbachia* about 6–46 mya (Meany *et al*. 2019).

#### *Wolbachia* divergence

Our chronogram analyses (Figure 3) estimate that *D. yakuba*-clade *Wolbachia* and three *w*Mel variants diverged about 29,000 years ago, and that *w*Tei split from sisters *w*San and *w*Yak about 2,500–4,500 years ago, with *w*San and *w*Yak diverging about 1,600-2,800 years ago. We estimate that the two most divergent *w*Mel variants from Richardson *et al*. (2012) and the reference *w*Mel genome split about 4,900–7,200 years ago, indicating more divergence among *w*Mel variants than among *D. yakuba*-clade *Wolbachia* strains. All of these results depend on the calibration provided by Richardson *et al*. (2012) and the relative accuracy of the underlying models of molecular evolution, which assume constant relative rates of change across data partitions. For the deepest divergence in Figure 3, between *D. yakuba*-clade *Wolbachia* and *w*Mel, we find that the estimated time depends on the variance in our prior distribution for substitution rates across branches, with a strict clock putting the divergence at about 73,000 years rather than 29,000 years obtained with the most variable prior. Despite this uncertainty, the quantitative differences of our *Wolbachia* divergence-time estimates do not alter the qualitative conclusion that these *Wolbachia* did not co-diverge with these hosts, which split several million years ago.

Our findings here and in Turelli *et al*. (2018) suggest that for several *Drosophila* species, their current *Wolbachia* infections have been in residence for only hundreds to tens of thousands of years. Bailly-Bechet *et al*. (2017) estimated *Wolbachia* residence times using data from more than 10,000 arthropod specimens spanning over 1,000 species. However, they analyzed DNA sequences from only part of the fast-evolving *fbpA Wolbachia* locus and the host *CO1* mtDNA locus. From an initial model-based meta-analysis, they concluded that “… most infections are very recent …” – consistent with our results. However, they also fit a more complex model with “short” and “long” time scales for acquisition and loss, conjecturing that short-term rates were associated with imperfect maternal transmission. Focusing on long-time rates, they concluded that *Wolbachia* infections persisted in lineages for approximately 7 million years on average, whereas lineages remained uninfected for about 9 million years. For *Drosophila*, such long infection durations would imply that *Wolbachia*-host associations often persist through speciation (see Coyne and Orr 1997 and Turelli *et al*. 2014 for estimates of speciation times in *Drosophila*, generally 10^5^–10^6^ years). Extrapolating from limited *Wolbachia* sequence data, Hamm *et al*. (2014) conjectured that cladogenic *Wolbachia* transmission might be common among *Drosophila*; but this extrapolation is refuted by our genomic analyses. The long *Wolbachia* durations proposed Bailly-Bechet *et al*. (2017) depend on their conjecture that their long-term rate estimates accurately reflect acquisition and loss of *Wolbachia* infections across species. This is worth testing with additional analyses of *Wolbachia*, mitochondrial and nuclear genomes from a broad range of arthropods.

#### *Wolbachia* acquisition––introgression versus horizontal

Our divergence-time estimates for the *D. yakuba*-clade *Wolbachia* versus their hosts preclude cladogenic acquisition. Unlike introgression, horizontal (or paternal) *Wolbachia* acquisition should produce discordance between phylogenies inferred for *Wolbachia* and the associated mitochondria. With the notable exception of *D. santomea* line Quija 37, which has mitochondria belonging to the clade associated with *D. yakuba*, but has *w*San *Wolbachia*, we find no evidence of discordance between the estimated mitochondrial and *Wolbachia* phylogenies. Hence our data indicate that acquisition by introgression is far more common than horizontal transmission between closely related species, consistent with data on acquisition of *w*Ri-like *Wolbachia* in the *D. melanogaster* species group (Turelli *et al*. 2018). Similarly, consistent with extensive data from *D. melanogaster* (Richardson *et al*. 2012) and *D. simulans* (Turelli *et al*. 2018) and smaller samples from *D. suzukii* and *D. ananassae* (Turelli *et al*. 2018), we find only one possible example of paternal transmission or horizontal transmission within *D. yakuba*-clade species.

We also investigated an alternative approach to distinguish between introgressive and horizontal *Wolbachia* acquisition by estimating substitution ratios for mtDNA versus *Wolbachia* (Turelli *et al*. 2018). Because we could estimate this ratio on each branch, we conjectured that this approach might have greater resolving power than our incompletely resolved mitochondrial and *Wolbachia* phylogenies. We expect higher ratios with horizontal transmission because mtDNA would have been diverging longer than recently transferred *Wolbachia*. However, this approach assumes that mtDNA and *Wolbachia* substitution rates remain relatively constant. This is contradicted by the finding of Ho *et al*. (2005) that mtDNA substitution rates decline substantially with increasing divergence time, reaching an asymptote after around 1–2 million years. To calibrate their *Wolbachia* substitution rate estimates, Richardson *et al*. (2012) used an experimentally observed mitochondrial mutation rate in *D. melanogaster* (6.2 × 10^−8^ mutations per third-position site per generation) that extrapolates to 62% third-position divergence per million years. This extrapolation is nonsensical as a long-term substitution rate. As summarized by Ho *et al*. (2005), typical mtDNA substitution rates are 0.5–1.5% per coding site per million years. Nevertheless, using the ratio of short-term rates for mtDNA and *Wolbachia*, Richardson *et al*. (2012) produced an estimate of the long-term *Wolbachia* substitution rate that agrees with independent estimates from *Nasonia* wasps (Raychoudhury *et al*. 2009) and *Nomada* bees (Gerth and Bleidorn 2016) derived from much longer divergence times (assuming cladogenic *Wolbachia* acquisition) (Conner *et al*. 2017). This paradox is resolved if *Wolbachia* molecular evolution is not subject to the dramatic slowdown in rates seen for mtDNA substitution rates.

The apparent difference between mtDNA molecular evolution (dramatic slowdown over longer time scales, Ho *et al*. 2005) and *Wolbachia* molecular evolution (relative constancy, as inferred from similar rates of differentiation over very different time scales) suggests why our relative-rate test does not reject introgressive transmission of *Wolbachia* between *D. melanogaster* and the *D. yakuba* clade, even though it is clearly impossible. A roughly 50-fold slowdown in mtDNA substitution rates over the time scale of the divergence of *D. melanogaster* from the *D. yakuba* clade, relative to the rate of differentiation within the *D. yakuba* clade, produces comparable mtDNA-*Wolbachia* substitution ratios for comparisons within the *D. yakuba* clade and between *D. melanogaster* and the *D. yakuba* clade. Because of this complication and our conjecture that relative rates of mtDNA versus *Wolbachia* substitutions over longer periods are likely to mirror the ten-fold differences we see for mtDNA versus nuclear genes, phylogenetic discordance between mitochondria and *Wolbachia* is clearly a much more robust indicator of horizontal (or paternal) *Wolbachia* transmission. Nevertheless, additional examples of cladogenic *Wolbachia* acquisition are needed to better understand relative rates and patterns of *Wolbachia*, mtDNA and nuclear differentiation over different time scales.

Our divergence-times estimate of the *D. yakuba*-clade *Wolbachia* versus their hosts precludes cladogenic transmission; and our phylogenetic analyses suggest that these species share very similar *Wolbachia* because of introgression, as originally argued by Lachaise *et al*. (2000). However, under either introgressive or horizontal transfer of *Wolbachia*, we expect the donor species *Wolbachia* sequences would appear paraphyletic when analyzed jointly with the *Wolbachia* from the recipient. Paraphyly allowed Turelli *et al*. (2018) to infer that *D. simulans* likely obtained its *Wolbachia* from *D. ananassae* and *D. subpulchrella* likely obtained its *Wolbachia* from *D. suzukii*. Paraphyly is generally expected soon after gene flow stops between populations. As noted by Hudson and Coyne (2002, Fig. 1), the time scale expected to produce reciprocal monophyly for mitochondria (and *Wolbachia*) under a neutral model of molecular evolution is on the order of the effective size of the species. Our results in Figure 3 indicate that, at least for our small samples, reciprocal monophyly for the *Wolbachia* in these three species has been achieved within a few thousand years. This suggests that reciprocal monophyly has been accelerated by species-specific selective sweeps within the *Wolbachia* or mitochondria of these species. This conjecture may be testable from estimates of host fitness using transinfected versus native *Wolbachia*.

### IS transposable elements mediate horizontal transfer of incompatibility loci between divergent *Wolbachia*

*Wolbachia* in all three *D. yakuba*-clade hosts cause both intra-and interspecific CI (Cooper *et al*. 2017), despite originally being characterized as non-CI causing (Zabalou *et al*. 2004; Charlat *et al*. 2004). CI is relatively weak, and its strength can vary among *w*Tei variants and *D. teissieri* backgrounds (Table 3 in Cooper *et al*. 2017). Differences in CI among *Wolbachia* variants has also been demonstrated in interspecific backgrounds where *w*Tei caused stronger CI in a *D. simulans* background (97.2 ± 1.3 SE percent embryo mortality) than either *w*Yak (26.5 ± 4.2 SE, percent embryo mortality) or *w*San (24.0 ± 4.1 SE, percent embryo mortality) (Zabalou et al. 2008). Both *w*Yak and *w*San induced CI in *D. simulans* comparable in intensity to that found by Cooper *et al*. (2017, Figure 3) in their original hosts. Surprisingly, Zabalou *et al*. (2008) found that the strength of CI induced by *w*Tei in *D. simulans* even eclipsed that of *w*Ri (89.8 ± 4.5 SE percent embryo mortality). These results must be reconciled with the fact that loci known to underlie CI do not vary within or among *D. yakuba*-clade *Wolbachia* variants we examined. The nearly complete CI induced by *w*Tei in *D. simulans* may depend on CI-causing factors yet to be identified or differences in gene expression.

In each *D. yakuba*-clade *Wolbachia* variant included in our analyses, we find a disruption of *cidB^w^*^Yak-clade^ with an inversion from amino acids 37–103 relative to the same region in sister *w*Mel. The inversion introduces multiple stop codons that could render this gene nonfunctional. Fixation of loss-of-function mutations in CI-causing loci is consistent with theoretical analyses showing that selection on *Wolbachia* within host lineages does not act to increase or maintain CI (Prout 1994; Turelli 1994; Haygood and Turelli 2009); indeed, we have also recently observed a single mutation that disrupts *cidB* in non-CI causing *w*Mau *Wolbachia* that infect *D. mauritiana* on the island of Mauritius (Meany et al. 2019). In both *w*Mau and the *D. yakuba*-clade *Wolbachia*, we find fixation of defects in the putative toxin gene. We expect that future genomic analyses will produce additional examples.

All *D. yakuba*-clade *Wolbachia* genomes included in our analysis harbor *cinA-cinB* loci originally discovered in the *w*Pip strain that diverged from A-group *Wolbachia*, including the *D. yakuba*-clade variants, about 6–46 mya (Meany *et al*. 2019). *cin* loci are also present in B-group *w*AlbB that infects *Ae. albopictus* and in A-group *w*NFe, *w*NPa, *w*NLeu, and *w*NFa *Wolbachia* that infect *Nomada* bees. *cin* loci are absent from *w*Mel, but the *w*Yak contig containing these loci is about 10% diverged from *w*Mel, while observed divergence between *w*Yak and *w*Mel across the rest of the genome is less than 1%. The *w*Yak-clade *cin* loci share about 97% similarity with the divergent B-group *w*Pip strain. *w*Yak-clade *cin* loci are more similar to *cinA-cinB* from the B-group *Wolbachia w*Pip and *w*AlbB than to those in A-group *Nomada Wolbachia* strains, which have two sets of *cin* loci that are as diverged from each other as they are from these regions in *w*Yak and in B-group *Wolbachia*. These observations suggest independent horizontal transfer of *cin* loci into *w*Yak and *Nomada Wolbachia*.

Our results indicate that independent of prophage movement, ISWpi1-element paralogs can move incompatibility loci via the excision of flanking ISWpi1 elements, followed by homologous recombination within the elements. Horizontal *Wolbachia* acquisition is common in *Drosophila* (Turelli *et al*. 2018) and other species (O’Neill *et al*. 1992), suggesting that double infections, which could provide the opportunity for ISWpi1-mediated transfer of incompatibility loci, may be common, even if transient. (A second infection need not become stably transmitted for horizontal gene transfer via ISWpi1 elements to occur.) In contrast, phage particles or virions could be introduced by a vector and provide the opportunity for ISWpi1 mediated transfer (Ahmed *et al*. 2015; Brown and Lloyd 2015), without the presence of a double *Wolbachia* infection. Determining whether the insertion of these loci was derived from a prophage region of the *Wolbachia* genome, or from a phage genome encapsulated in a phage particle, remains an open question. While the ISWpi1 element in *w*Mel (Mel #9 labeled “B” in Figure 5; Cordaux 2008) is not part of our *w*Yak assembly, homologs of this element are present in assemblies of several other A-group *Wolbachia* including *w*Inc and *w*Ri (Cordaux *et al*. 2008; Bordenstein and Bordenstein 2016). We predict this element occurs in the unassembled region of our *w*Yak assembly. Footprints of ISWpi1 elements in the region flanking the *cinA* genes for both copies of the gene in the *Nomada Wolbachia* provide further support for our hypothesis. Long-read-based *Wolbachia* assemblies from many infected host systems will elucidate the role of ISWpi1 elements in horizontal transfer of CI loci. Overall, the ecology of horizontal *Wolbachia* transmission is crucial to understanding *Wolbachia* acquisition; and the transfer and dynamics of CI loci are crucial to understanding *Wolbachia* evolution.

## SUPPLEMENTAL INFORMATION

### SUPPLEMENTAL MATERIALS AND METHODS

#### PCR amplification and Sanger sequencing of “A”, “B”, and “C” regions in Figure 5

DNA was extracted from each isofemale (yakuba_S09_L24, yakuba_S01_L19, and yakuba_S01_L31) line using a standard ‘squish’ buffer protocol (Gloor *et al*. 1993). A standard PCR assay (Simpliamp ThermoCycler, Applied Biosystems, Singapore) was performed using the GoTaq™ DNA Polymerase (Promega™, Wisconsin, USA) master mix. PCR program began with 3 minutes at 94°C, followed by 34 cycles of 30 seconds at 94°C, 30 seconds at 55°C, and 1 minute and 15 seconds at 72°C. The profile finished with 8 minutes at 72°C. The amplified PCR products were visualized on a 1% agarose gel that included a ladder.

We evaluated regions A, B, and C in Figure 5 using PCR and Sanger sequencing. This includes one primer set covering the A region, four primer sets attempting to cover the B region, and one primer set covering the C region (Table S1). We performed touchdown PCR with an annealing gradient from 70°C to 55°C, which provided the same results as the standard PCR conditions. In cases where no products were returned, we lowered the annealing temperature from 55°C to 50°C and repeated the PCR. PCR products from a focal genotype (yakuba_S09_L24) were cleaned using the QiaQuick PCR purification kit (Qiagen, Germany). We sent the cleaned PCR products to Eurofins (Kentucky) for Sanger sequencing. The loci, primers, and product lengths of PCR experiments are presented in Table S1.

### SUPPLEMENTAL TABLES

**Table S1.**
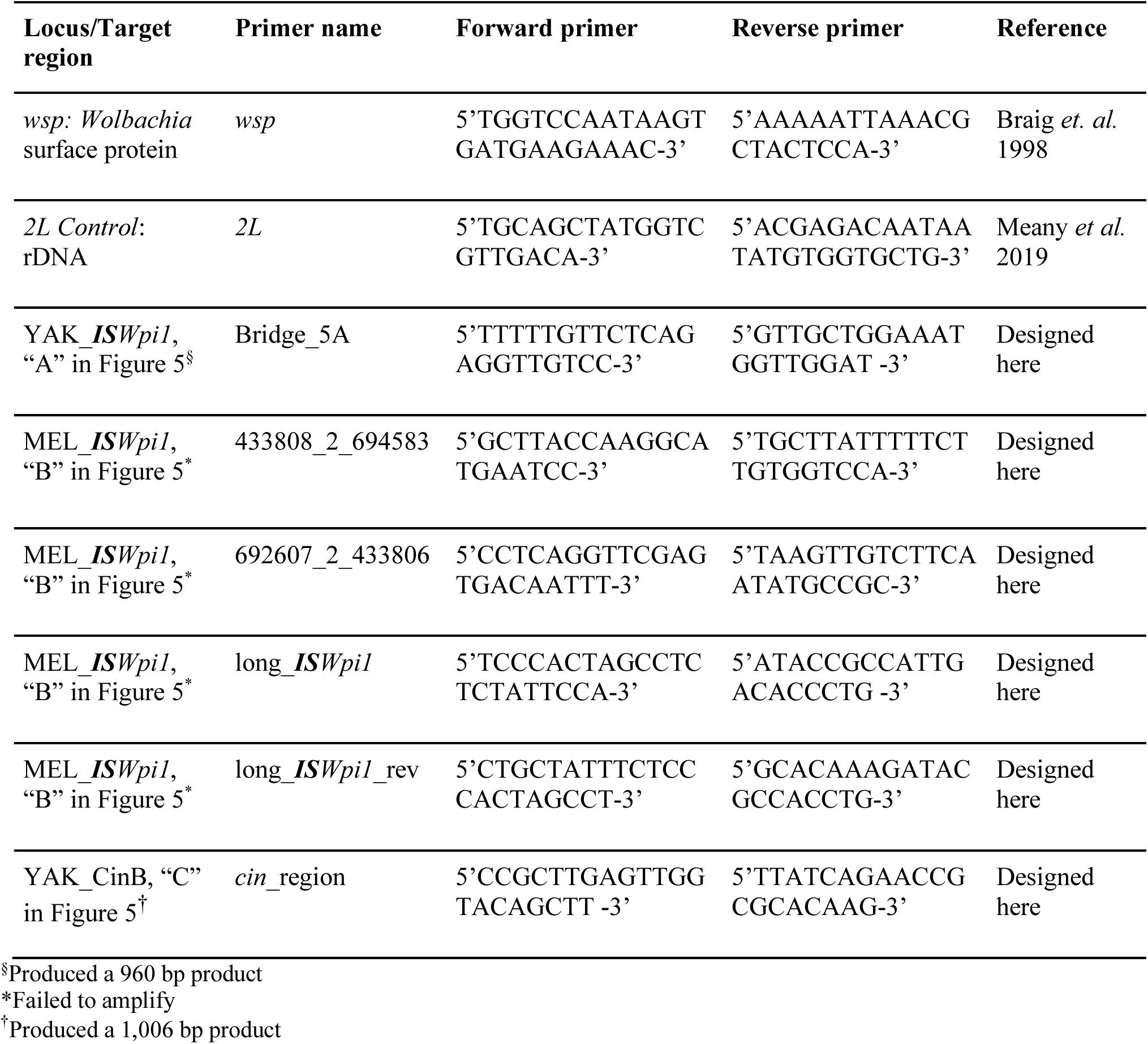
Primers used in this study.

**Table S2.** The scaffold count, N50, and total assembly size of each *Wolbachia* pseudoreference.

| Genome | Scaffold count | N50 | Total assembly size |
| --- | --- | --- | --- |
| wYak | 118 | 17888 | 1.28 million bases |
| wSan | 300 | 7231 | 1.18 million bases |
| wTei | 122 | 16157 | 1.30 million bases |

**Table S3.** Near-universal, single-copy proteobacteria genes used to assess assembly completeness. We used BUSCO v. 3.0.0 to identify complete and fragmented proteobacteria genes (out of 221) found in our draft *Wolbachia* genomes. For comparison, we provide results for reference-quality *Wolbachia* genomes.

| Genome | Complete | Fragment | Not found |
| --- | --- | --- | --- |
| **Complete reference genomes** |  |  |  |
| wRi | 180 | 2 | 39 |
| wMel | 180 | 2 | 39 |
| wAu | 181 | 2 | 38 |
| wHa | 179 | 3 | 39 |
| wNo | 181 | 4 | 36 |
| **Draft genome assemblies** |  |  |  |
| wYak | 181 | 2 | 38 |
| wSan | 181 | 2 | 38 |
| wTei | 180 | 2 | 39 |

**Table S4.** Rate multipliers for the *Wolbachia* relaxed-clock chronograms. All numbers are estimated from the Γ(2,2) trees. They are the total estimated substitutions per site for that partition across the relevant clade (See Methods).

|  | 1<sup>st</sup> position<br>(95% credible interval) | 2<sup>nd</sup> position<br>(95% credible interval) | 3<sup>rd</sup> position<br>(95% credible interval) |
| --- | --- | --- | --- |
| *D. yakuba*-*Wolbachia* | $1.9 \times 10^{-5}$,<br>($9.6 \times 10^{-6}$, $3.0 \times 10^{-5}$) | $1.2 \times 10^{-5}$,<br>($5.2 \times 10^{-6}$, $1.9 \times 10^{-5}$) | $1.7 \times 10^{-5}$,<br>($8.5 \times 10^{-6}$, $2.7 \times 10^{-5}$) |
| wMel | $4.4 \times 10^{-5}$,<br>($2.2 \times 10^{-5}$, $7.1 \times 10^{-5}$) | $5.7 \times 10^{-5}$,<br>($2.8 \times 10^{-5}$, $9.2 \times 10^{-5}$) | $3.4 \times 10^{-5}$,<br>($1.5 \times 10^{-5}$, $5.6 \times 10^{-5}$) |
| *D. yakuba*- clade and<br>wMel *Wolbachia* | $1.7 \times 10^{-4}$,<br>($1.1 \times 10^{-4}$, $2.4 \times 10^{-4}$) | $1.4 \times 10^{-4}$,<br>($8.7 \times 10^{-5}$, $1.9 \times 10^{-4}$) | $2.0 \times 10^{-4}$,<br>($1.2 \times 10^{-4}$, $2.7 \times 10^{-4}$) |

**Table S5.** Divergence estimates and credible intervals for the *Wolbachia* relaxed-clock chronograms estimated from the Γ(7,7) branch-rate prior.

| **Relaxed-clock analysis** | **wMel-wYak divergence** | **wYak-clade divergence** | **wMel-clade divergence** | **wYak-wSan divergence** |
| --- | --- | --- | --- | --- |
| *D. yakuba-Wolbachia* | NA | 2642,<br>(781, 5904) | NA | 1545,<br>(401, 3680) |
| wMel | NA | NA | 4955,<br>(1363, 11093) | NA |
| *D. yakuba*- clade and<br>wMel *Wolbachia* | 49585,<br>(17452, 106310) | 3914,<br>(1432, 9024) | 7908,<br>(2511, 18739) | 2252,<br>(740, 5229) |

### SUPPLEMENTAL FIGURES

**Figure S1.**
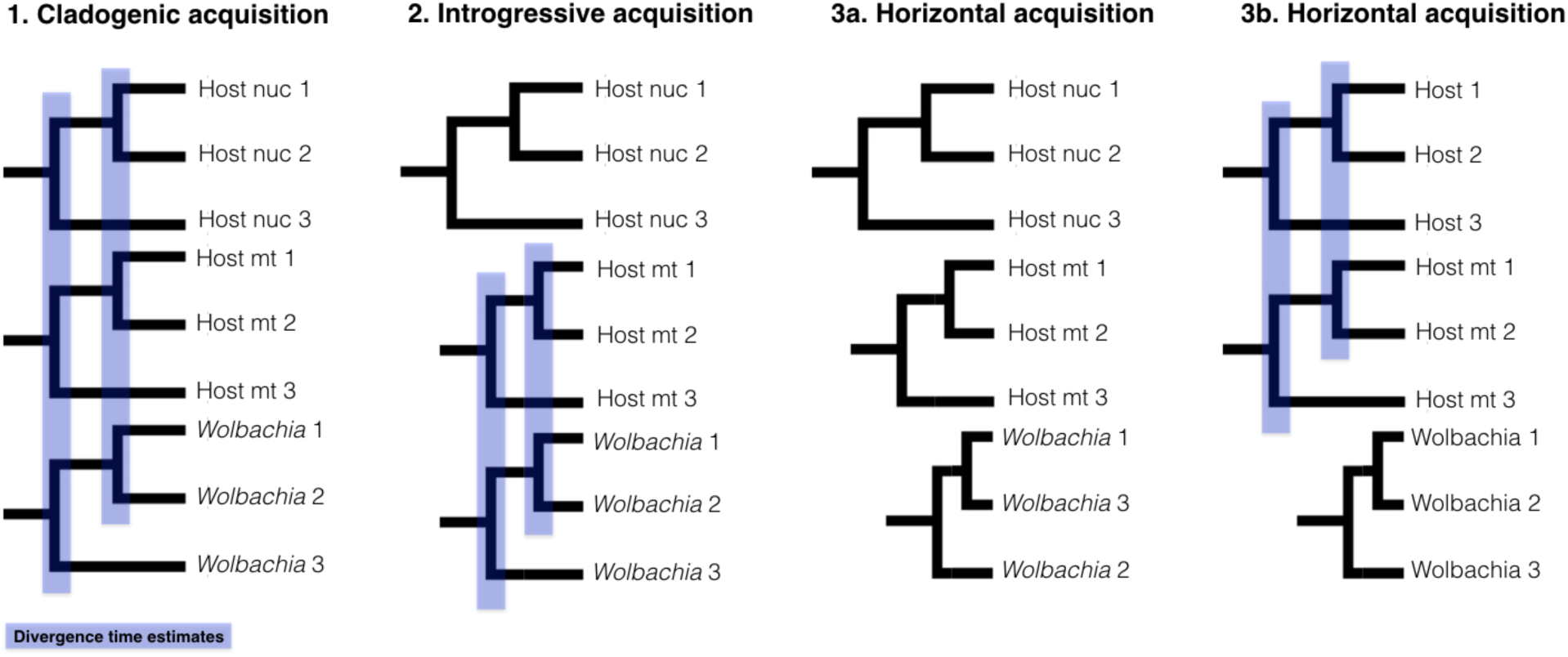
Phylogenetic hypotheses about *Wolbachia* acquisition. Infections could be 1) old and codiverging with host genomes (cladogenic acquisition), 2) codiverging with mtDNA (mt) with both mtDNA and *Wolbachia* having more recent divergence times than host nuclear genomes due to a history of host hybridization (introgressive acquisition), or 3) relatively young with more recent divergence times than host genomes (horizontal acquisition). For introgressive acquisition, mitochondrial and *Wolbachia* topologies could be discordant with the host nuclear topology. For horizontal acquisition, *Wolbachia* genomes could be discordant (3a) or concordant (3b) with host genomes. *Wolbachia* could be acquired horizontally by hosts with a history of hybridization introgression (3a) or by hosts with no history of hybridization and introgression (3b). Note that *Wolbachia* topologies could be concordant or discordant with host genomes in either case of horizontal acquisition.

**Figure S2.**
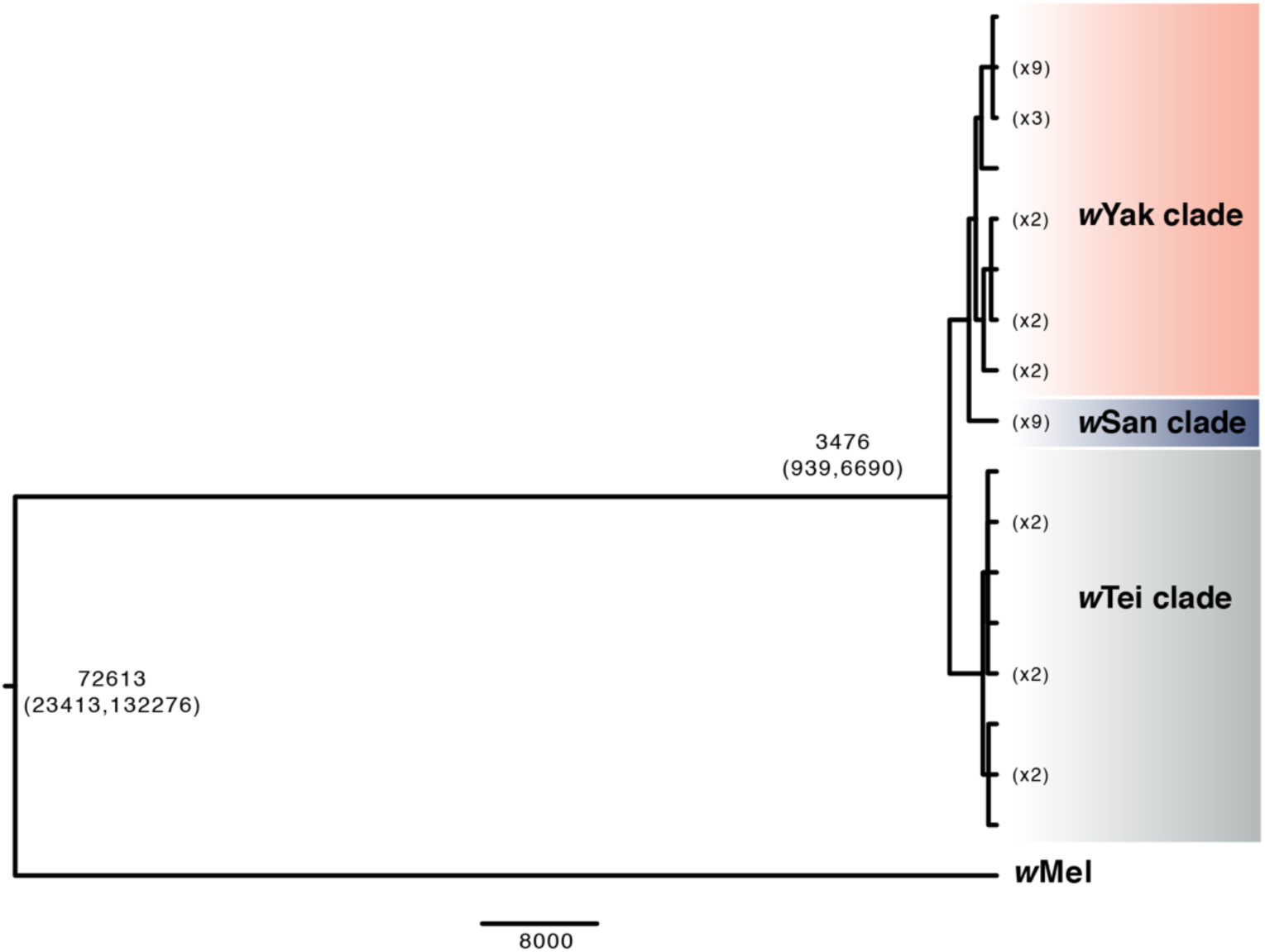
A Bayesian strict-clock *Wolbachia* absolute chronogram using 621 genes containing a total of 624,438 bp. Nodes with posterior probability less than 0.95 were collapsed into polytomies. Key nodes are labeled with age estimates.

**Figure S3.**
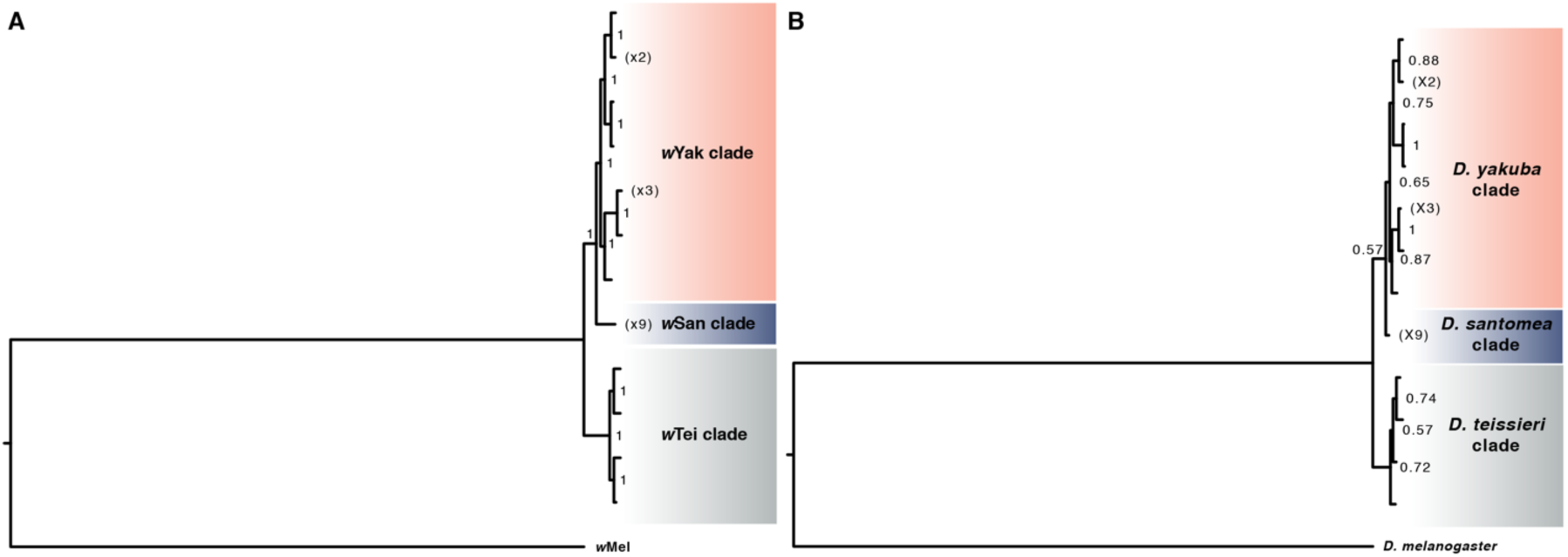
A. Bayesian *Wolbachia* phylogram using 621 genes containing a total of 624,438 bp. B. Bayesian mtDNA chronogram using the thirteen protein-coding genes. Nodes are labeled with their posterior probability.

**Figure S4.**
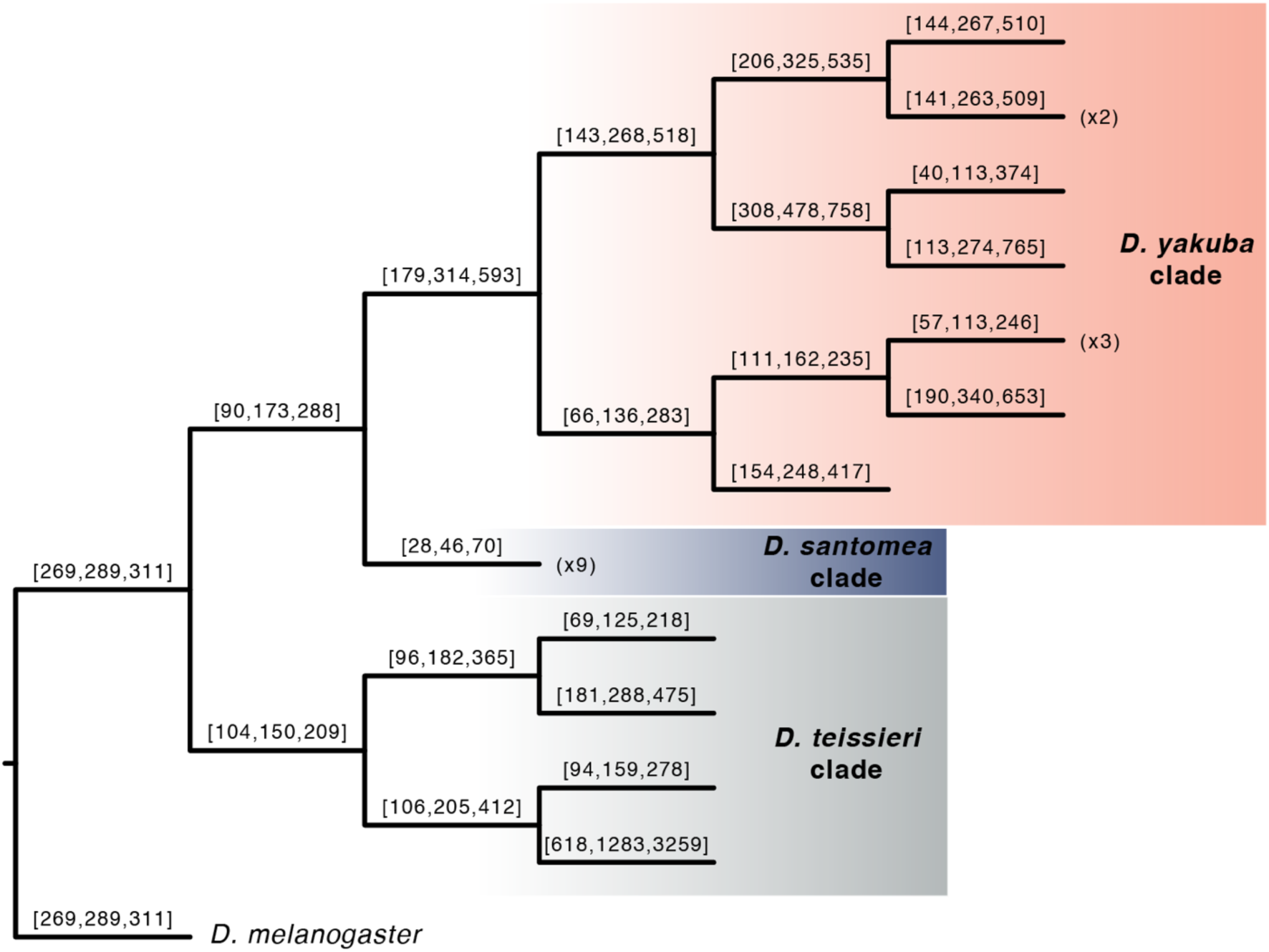
A Bayesian phylogram with all branches fixed to length 1. The topology is the consensus of the mitochondrial and *Wolbachia* phylograms. Branches are labeled with the quartiles of the posterior distribution of ratios of the substitution rates of mtDNA versus *Wolbachia* along the branch (see Methods).

